# Sequencing of historical plastid genomes reveal exceptional genetic diversity in early domesticated rye plants

**DOI:** 10.1101/2024.08.08.607158

**Authors:** Jovan Komluski, Sofia Filatova, Frank Schlütz, Benjamin Claaßen, Manfred Rösch, Ben Krause-Kyora, Wiebke Kirleis, Eva H. Stukenbrock

**Affiliations:** Botanical Institute, Kiel University, 241148 Kiel, Germany; Max Planck Institute for Evolutionary Biology, 24306 Plön, Germany; Groningen Institute of Archaeology, Groningen, the Netherlands; Institute for Prehistoric and Protohistoric Archaeology, Kiel University, 24118 Kiel, Germany; Archaeobotanical Laboratory of the State Office for Cultural Heritage Baden Württemberg, 78343 Hemmenhofen/Gaienhofen, Germany; Institute of Clinical Molecular Biology, Kiel University, 24118 Kiel, Germany

**Keywords:** Ancient DNA, *Secale cereale*, evolution, chloroplast sequences, breeding, diversity

## Abstract

In medieval central Europe, rye was one of the most important agricultural crops. It’s properties of frost resistance, general resilience and resistance to many pathogens made it invaluable for medieval farmers. Rye has a distinct domestication history compared to other cereal crops and was not domesticated directly from its wild ancestors, like barley and wheat. Rye is considered to be a “secondary domesticate”, i.e. a crop with domestication traits that initially evolved as an arable weed but eventually was intentionally sown and propagated as a crop. To study the history of rye domestication, genetic sequences of present-day plant populations as well as material from historical samples can provide insights into the temporal and spatial signatures of domestication. In this study we combined archaeobotanical methods and ancient DNA sequencing of well-preserved, historical rye material to study patterns of genetic diversity across four centuries. We first applied archaeobotanical methods to characterize rye material acquired from construction material ranging from the 14^th^ to 18^th^ century from different locations in Germany. Next, we extracted DNA to sequence complete chloroplast genomes of six individual samples. We compared the 115,000 bp chloroplast genomes of historical rye samples to chloroplast genomes of other cereal crops and identified 217 single nucleotide variants exclusive to historical samples. By comparing the aDNA chloroplast samples with modern rye chloroplasts, we show that the genetic variation in ancient rye populations was exceptionally high compared to samples from contemporary rye cultivars. This confirms that late domestication and selective breeding have reduced genetic variation in this important crop species only in the last few centuries.

**Highlights:**

- Historical plant material covering four centuries was obtained from half-timbered houses from five locations in Germany
- Integrative archaeobotanical analyses and ancient DNA sequencing provided insights into genetic diversity of rye plants from historical farmland fields.
- Sequence analyses of complete assembled chloroplast genomes reveal expectational diversity in rye populations.
- Late domestication of rye preserved genetic diversity over centuries. The more recent intensification of rye breeding has however conferred a considerable loss of genetic diversity in this important crop.

## 1. Introduction

Rye, *Secale cereale,* is an important crop in eastern, central and northern Europe. It is produced in other regions including North and South America and China although to a far lesser extent than in Europe. Rye is mainly cultivated for bread production, for beer brewing and whiskey distillation, as well as for animal fodder. The rye plant thrives in nutrient poor soils, can resist frost and is known to be resistant to many fungal diseases making it a valuable agricultural crop (Persson and Bothmer, 2022; Schlegel, 2014).

The evolutionary history of rye has been much disputed, but ample evidence suggests that rye was domesticated from weedy rye species rather from wild species (Schreiber et al., 2021). Wild *S. cereale* relatives occur in the “Fertile Crescent” region along with wild relatives of wheat and barley. This indicates that the three cereal species share the same center of origin. However, archeological remains of rye, from the pre-pottery Neolithic, are always found along with remains of either barley or wheat.

*Secale cereale* ssp. *vavilovii* is considered the ancestor of domesticated rye, occurring in parts of Armenia and Eastern Turkey (Hagenblad et al., 2016; Skuza et al., 2019; Zohary et al., 2012). The domesticated form of rye first appeared during the seventh millennium BC as an admixture in stores of domesticated einkorn (*Triticum monococcum* ssp. *monococcum*) and emmer (*Triticum turgidum* ssp. *dicoccum*) from South Central Anatolia and the Upper Euphrates valley in Syria (Hillman, 1975; Nesbitt, 2002), suggesting that it developed traits of domestication while growing as an arable weed (Hillman, 1975). Intriguingly, recent studies have indicated the gathering or cultivation of a perennial species of wild rye (*Secale montanum*) as early as the Natufian (ca. 11,100-10,000 BC) in the Middle Euphrates area (Douché and Willcox, 2023). The earliest evidence of cultivation targeted at domesticated rye dates back to the fourth and second millennium BC in Central Anatolia, but based on current archaeobotanical evidence it was most likely a sporadic practice that did not yet establish itself as firmly as the cultivation of barley and species of wheat (Hillman, 1978; Zohary et al., 2012).

After having entered Europe as an arable weed around the fifth millennium BC and reaching areas north of the Alps, rye retained its status as an arable weed for the next thousands of years (Behre, 1992). Occasional archaeobotanical finds of rye grains as weedy admixtures have been found to increase in numbers from 2000 BC to 400 AD, within the last two millennia, gradually developing into one of the main staple crops in medieval Europe (Behre, 1992; Golea et al., 2023). The turn of rye from a weed into a domesticated crop in the course of the Iron Age may have been favored by colder weather conditions, a permanent field culture and a change in harvest practices with the introduction of iron tools. Since it was not domesticated directly from its wild ancestors, like barley and wheat, rye is attributed a “secondary domesticate”, i.e. a plant with domestication traits that is initially considered an arable weed but eventually develops into an intentionally sown crop in various regions independently, outside of its area of origin (Barret, 1983; Hawkes, 1969; Hillman, 1981). First archaeobotanical assemblages dominated by rye are known from northern-central Europe from about the second century AD onward and prove the development and use of rye as a crop (Behre, 2010, 1992; Grikpėdis and Matuzevičiūtė, 2016).

In the 12^th^-13^th^ century rye was one of the major crops grown in Europe, likely favored by its “robustness” and tolerance to unfavorable environmental conditions. As documented by pollen records, rye became the main cereal throughout most of northern and central Germany around 1000 AD and it was not until the middle of the 20th century that rye was replaced by bread wheat in its dominant position (Miedaner, 2014). Due to cross-pollination and the transportation of pollen by the wind over long distances (Behre, 1981), it took until the middle of the 19th century for first successful targeted selection of varieties from landraces like the Probstei rye. Around 1880, intensified breeding efforts led to rye varieties with greater resistance to lodging (Schindler, 1920; Schlegel, 2014), and a 100 years later, around 1980, hybrid breeding resulted in a considerable increase in yield, but also in greater genetic uniformity, especially uniformity of the cytoplasm (Miedaner 2014, Schlegel 2014). Today, some wild rye taxa are used as gene sources for crop improvement by interspecific and intergeneric crosses with barley and wheat (Schlegel 2014).

Genetic analyses of rye have been greatly facilitated by the availability of a reference genome sequence (Li et al., 2021). The exceptionally large genome of rye, approximately 7.8 Gb, has been used to map genetic variants of different rye cultivars, land races and wild sister species (Sun et al., 2022). By including whole genome data from the weedy rye species *S. cereale* subsp. *vavilovii* and from the wild rye species, *S. strictum* and *S. sylvestris*, Li and colleagues demonstrated that domesticated rye mostly likely derived from the weed *S. cereale* subsp. *vavilovii* supporting previous conclusions that have been based on archeological remains (Li et al., 2021).

In the medieval Europe, rye was harvested for the grains, but the up to 2-meter-tall cereal crop has been used for other purposes (Behre, 1992). Half-timbered houses have been built in Europe for centuries, and some well-preserved buildings dating back to medieval times can still be found. Infill material of these constructions generally comprised a mixture of clay and chalk with a binder such as grass or straw and water. Straws of rye have been popular as infill material in half-timbered houses, and can, today, be found inside well-preserved constructions. Such remains of rye and other straw material in half-timbered houses from medieval times, provide a unique opportunity to investigate the diversity of rye as it was centuries ago. In this study we explored the access to so-called “*Wellehölzer*” from half-timbered houses collected at different sites in Germany and representing different epochs ranging from the 14^th^ to the 19^th^ century.

In this study, we make use of ancient DNA sequencing approaches to reconstruct chloroplast genomes from different rye individuals and we combine comparative genetic analyses with archaeobotanical descriptions of the well-preserved plant remains. We demonstrate extensive genetic variation in the rye chloroplast DNA, including variable sites in protein coding genes. Moreover, we show that the composition of plant remains suggest that rye was sown in highly disturbed fields and likely cultivated as a winter cereal. Our study points to the great potential of ancient DNA analyses and archaeobotanical investigations to unravel the composition and history of agricultural crops.

## 2. Materials and methods

### 2.1. Plant material

Plant material used in this study consisted of 17 samples of historical plant material from construction parts of half-timbered houses (plant-tempered clay, with focus on *Wellerhölzer*) built between the 14^th^ and 19^th^ century from five locations in Germany (**Fig. 1**). Seven *Wellerhölzer* samples were collected from the old town hall in Göttingen, and the remaining ten historical samples were obtained from the south German half-timbered houses in Schwäbisch Hall (five samples), Reutlingen (three samples), Zaisenhausen/Mulfingen (one sample) and Weipertshofen (one sample) (**Fig. 1**). According to available historical data, for the buildings from which the material was collected, the plant material isolated from these locations, dates from 14th to 19th century according to available historical data for the buildings from which the material was collected (Fischer and Rösch, 2008; Rösch and Fischer, 2001, 1999). We note that radiocarbon dates taken on material dating between 1650 and 1950 AD unfortunately cannot be established with certainty due to the Maunder Minimum and the Suess effect, which greatly impacted the 14C content in the atmosphere (Keeling et al., 2017; Rek, 2010). Radiocarbon dates falling within the scope of these atmospheric events results in multiple possibilities and can thus only be indicated broadly. In this study, we therefore limit ourselves to the construction dates of the buildings, being aware that the actual dates of the plant remains may be younger due to later reconstruction activities. A table listing the samples, their origin and 14C dates is provided in **Tab. S1**.

**Fig. 1.**
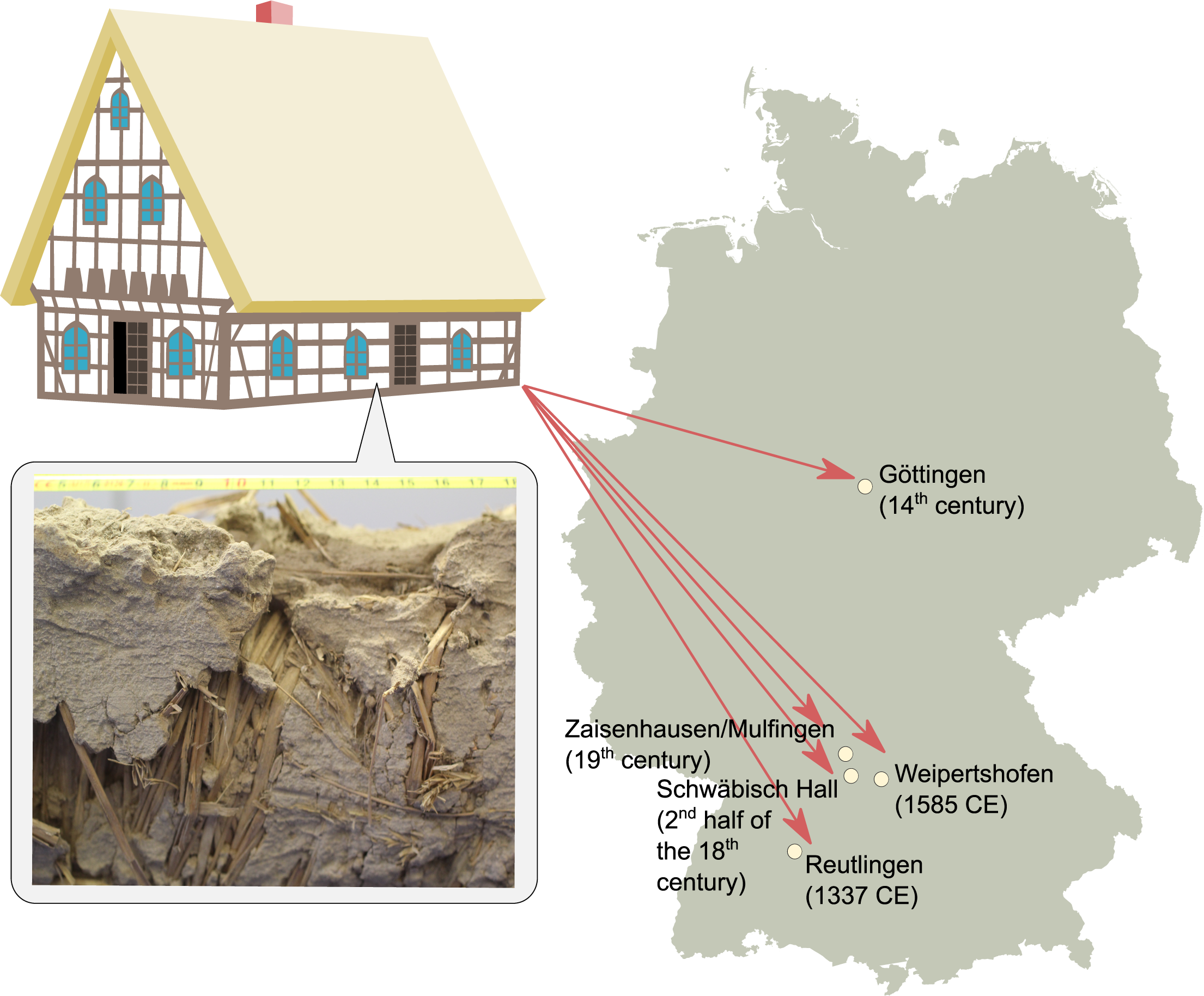
Sampling sites and ages of *Wellerhölzer* plant material used in the study. Historical rye varieties were collected from construction material of traditional half-timbered German houses from five locations designated on the map.

### 2.2. Archaeobotanical analysis

Archaeobotanical analyses of the plant material was performed by different individuals at different occasions. The material from Schwäbisch Hall, Reutlingen and Zaisenhausen/Mulfingen was analysed by Rösch and Fischer (Fischer and Rösch, 2008, 1999; Rösch and Fischer, 1999, 1997), the material from Weipertshofen by Fischer and the material from Göttingen (Filatova et al., 2021). The plant remains were separated from the sediment matrix by soaking in water and subsequently sieving the material using (stacked) sieves with mesh widths varying from 2 mm to 0.25 mm. The remains were dried before being analyzed using low-magnification (up to 40 x) stereo microscopes. Cultivation practices and soil conditions of the rye used in the two *Wellerhölzer* from Göttingen and Schwäbisch Hall were reconstructed based on a combination of autecology and functional autecology of the presumed arable weed taxa (for a more elaborate discussion, see (Filatova et al., 2021)). These two approaches are based on the notion that characteristics of plants are indicative of certain climatic conditions (e.g., temperature and light), soil conditions (e.g., pH and nitrogen), and activities of soil disturbance (e.g., digging and trampling) (Bogaard et al., 1999; Charles et al., 2002; Duckworth et al., 2000; Ellenberg, 1988; Fischer and Rösch, 2008).

Taxa that were identified to species level were included in the analysis. Functional attributes and autecological characteristics of the selected species were used to reconstruct cultivation practices (**Appendix 1**, **Tab. S2**). Data on synecology, autecology, and functional attributes were collected from (Ellenberg, 1988; Klotz and Kühn, 2002; Kühn et al., 2004).

### 2.3. Sample preparation, DNA extraction and sequencing

The ancient plant material used in this study included straw, spikelets, and ears of cereals which were identified to belong to rye as well as wheat and barley based on visual examination of morphological traits (**Tab. S1**). To prepare selected rye samples for the DNA extraction, approximately 0.1–0.2 g of desiccated rye material of each sample was washed in a 2% sodium hypochlorite (NaOCl) solution, followed by an additional ethanol (C_2_H_5_OH) wash for the surface cleaning from contaminants.

Ancient DNA was extracted according to a previously published protocol specifically developed for the extraction of ancient DNA using N-phenacyl thiazolium bromide (PTB) with minor modifications (Wales and Kistler, 2019). In short, the washed plant remains were frozen with dry ice and grinded, overnight incubation at 37 °C in a buffer containing N-phenacyl thiazolium bromide (2.5 mM PTB, 50 mM dithiothreitol (DTT), 10 mM TRIS, 10 mM ethylenediaminetetraacetic acid (EDTA), 5 mM sodium chloride (NaCl), 1% sodium dodecyl sulfate (SDS), 0.4 mg/ml Proteinase K) in a rotator. After incubation, DNA was extracted from and converted into partial Uracil-DNA glycosylase (UDG) libraries (Rohland et al., 2015)17, following the established laboratory guidelines for aDNA work (Cooper and Poinar, 2000)17.

Sequencing was performed on the Illumina NovaSeq6000 platform at the Institute of Clinical Molecular Biology, Kiel University, Germany using the NovaSeq S1 Kits with 200 cycles.

Illumina reads were deposited at https://doi.org/10.5281/zenodo.12655083.

### 2.4. Read mapping, variant calling and annotation

In our study, we have focused on the chloroplast genome that occurs in higher copy numbers than the nuclear genome, which congruently is reflected in sequence abundance. To assess the nucleotide diversity of the chloroplast genome between the modern and ancient rye, Illumina paired reads from *Wellerhölzer* rye samples were mapped to the KC912691 reference chloroplast genome of rye (Middleton et al., 2014) with the nf-core/EAGER, bioinformatic pipeline specifically tailored for the computational analysis of ancient DNA (Peltzer et al., 2016). The same approach for read mapping and variant calling was conducted for the *Wellerhölzer* wheat (KP210077), barley (KP210078) and Cerealia (KP210082, KP210083, KP210084, KP210086) samples that also we also included here in our phylogenetic analysis. To this end, reads from *Wellerhölzer* wheat samples were mapped to the KC912694 reference chloroplast genome of wheat, and reads from *Wellerhölzer* barley were mapped to the KC912687 reference chloroplast genome of barley (Middleton et al., 2014), while the *Wellerhölzer* Cerealia samples were mapped to the reference chloroplast genome of rye.

We assessed the aDNA damage mapping profiles via the mapDamage tool incorporated in the nf-core/EAGER pipeline (Peltzer et al., 2016), in order to detect specific nucleotide misincorporations and DNA fragmentation signatures in sequencing reads, including an increase of C→T and G→A substitutions at the 5’ and 3’ regions of the sequenced DNA, a typical signature in degenerated ancient DNA (Ginolhac et al., 2011). Mapped reads from which duplicates were removed by the EAGER pipeline and which otherwise passed the mapping-damage-control and quality check, were further used for the variant calling. Genetic variants from the chloroplast genome of medieval cereals were called by samtools mpileup (version 1.7, options -C 50 -Q 0 -q 20 -uf), followed by bcftools call (version 1.7, options -vc - O u -o) (Danecek et al., 2021). For our analyses, we have focused exclusively on biallelic SNPs. Therefore, called variants were filtered for biallelic SNPs with bcftools view, and only SNPs with mapping quality higher than 20 were used for the downstream analyses.

To perform functional annotation of the SNPs identified in the *Wellerhölzer* rye chloroplast genome, vcf (variant call format) files from all *Wellerhölzer* rye samples were merged into a single vcf file via vcf-merge program from vcftools (version 0.1.15) (Danecek et al., 2011). The merged vcf file was further used as an input for SnpEff (version 5.1d), a program for annotating and predicting the SNP effects (Cingolani et al., 2012). The number of alleles with a specific annotation was calculated from the output SnpEff file with a custom-made Python script. Lastly, the number of SNPs per specific chloroplast gene was calculated with a custom Python script from an output file from bedtools intersect (v2.26.0, options -wa -wb) (Quinlan and Hall, 2010) between the gene annotation file of the rye reference chloroplast genome and bed file with the coordinates of SNPs identified in *Wellerhölzer* rye samples. The SNP number per chloroplast gene was normalized for gene size and calculated per 1000 bp windows with a custom-made Python script. The custom-made scripts used for the analyses and the plotting of figures are provided in **Appendix 2**.

### 2.5. Chloroplast sequence variation and phylogenetic analysis

To characterize the chloroplast sequence variation between the *Wellerhölzer* samples, we identified all genetic variants and distinguished SNPs specific to certain sampling sites or shared across different sampling sites (**Fig. 2A** and **Fig. S1**). Further, for phylogenetic analysis, we generated a Neighbor-Joining tree that included chloroplast sequences of the historical *Wellerhölzer* samples, as well as the chloroplast sequences of modern rye varieties, rye sister species and reference chloroplast sequences of wheat and barley (**Fig. 2B**). For the rye species, five chloroplast sequences of modern rye varieties (Bernhardt et al., 2017), and eight chloroplast sequences from five additional rye sister species (Bernhardt et al., 2017; Ding et al., 2022; Du et al., 2022; Skuza et al., 2022) were included in the Neighbor-Joining analysis, as well as the reference chloroplast sequences of *Triticum aestivum* and *Hordeum vulgare* (Schreiber et al., 2020; The International Wheat Genome Sequencing Consortium (IWGSC) et al., 2018).

**Fig. 2.**
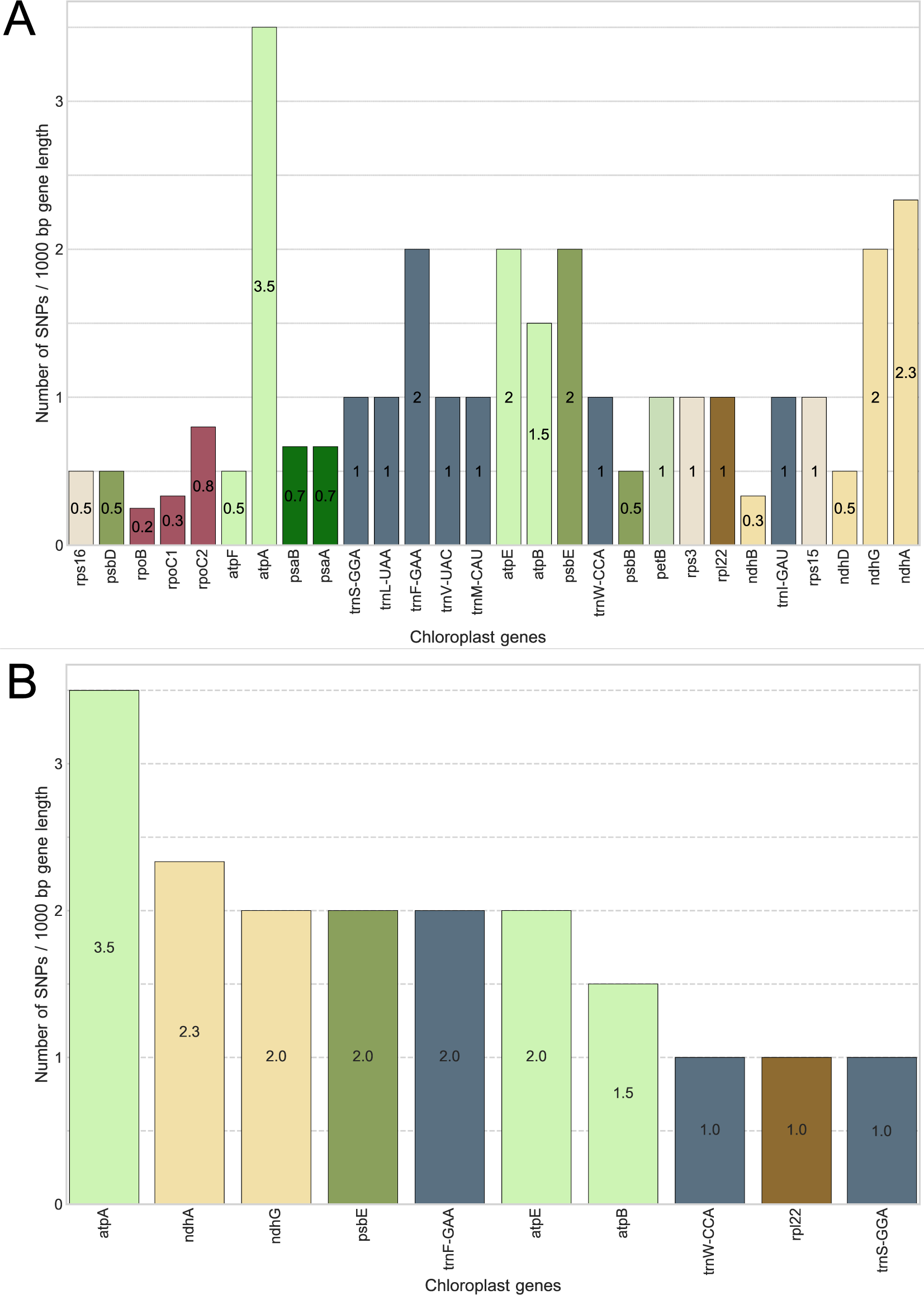
Chloroplast sequence variability in the ancient *Wellerhölzer* rye. **A)** Upset plot showing intersections between ancient SNPs in the *Wellerhölzer* rye samples grouped based on their sampling location. **B)** Neighbor-Joining tree displaying the genetic relationship between the chloroplast sequences of the historical *Wellerhölzer* cereals and modern cereal accessions. Branch node colors designate the chloroplast sequences of the respective cereal species in accord to the legend. The numbers above branches represent the branch support value.

For the chloroplast sequences of the historical *Wellerhölzer* rye and Cerealia samples, we generated consensus sequences using the reference rye chloroplast sequence KC912691 (Middleton et al., 2014) and the vcf file of a respective historical sample via bcftools consensus (Danecek et al., 2021). For the two chloroplast sequences of wheat and barley, a consensus sequence was generated with the reference chloroplast sequence of wheat (Middleton et al. 2014) and barley (Middleton et al. 2014), respectively. Multiple sequence alignments between were constructed using ClustalW (Thompson et al., 1994) which were subsequently used to generate a Neighbor-Joining tree (using 1000 bootstraps) with the SeaView software (Gouy et al., 2010). Trees were visualized using Figtree (v1.4.4) (http://tree.bio.ed.ac.uk/software/figtree/).

## 3. Results

The results presented here encompass archaeobotanical and molecular analysis of plant remains from half-timbered houses (*Wellerhölzer*) from five locations in central and south Germany dating from the 14th to 19th century AD. Previous archaeobotanical analyses were conducted on thousands of remains from *Wellerhölzer*, and were combined here with molecular analyses focusing on the chloroplast sequences of plant remains identified as rye.

### 3.1. Archaeobotanical analyses reveal patterns of crop rotation

The individual archaeobotanical analyses have shed light on the conditions in which the rye grew, including soil pH, the season of cultivation and, for a part, the likely cultivation methods. Based on ecological characteristics of the arable weeds accompanying the rye, it was defined that rye was cultivated on moderately basic soils in Zaisenhausen/Mulfingen (Fischer and Rösch, 2008) and Göttingen (Filatova et al., 2021), while it grew on more acidic soils in Weipertshofen (Rösch and Fischer, 2001), Reutlingen (Rösch and Fischer, 1999) and Schwäbisch Hall (Appendix 1; Tab. S2) (Rösch and Fischer, 1997). In all cases, the seasonality of the arable weeds indicates that the rye was sown in autumn as a winter cereal. The crop spectra uncovered from the investigated construction components indicate that it is likely that winter rye was grown in rotation with summer crops such as oat (*Avena sativa*) and barley (*Hordeum vulgare*), possibly in a three-field system. In Göttingen, the analysis furthermore indicated that the soil was likely manured and worked with a moldboard plough. Taken together, archaeobotanical analyses suggest that, aside from variation in soil pH, rye was cultivated using similar methods and in the same season across all the investigated locations in Germany. The variation in soil pH can be explained by the type of soils available for crop cultivation in the respective regions as well as by the addition of manure (Filatova et al., 2021; Fischer and Rösch, 1999).

### 3.2. Exceptional variation in *Wellerhölzer* rye chloroplast genomes

To assess the overall genetic variation among the eleven *Wellerhölzer* rye samples, we first identified single nucleotide polymorphisms (SNPs) in the chloroplast genomes and grouped SNP intersections based on the sampling site (**Fig 2A**). 45% of the identified SNPs (97 out of 217) were unique to the sampling site (i.e., variants at certain chloroplast genome positions occurred only in samples from a specific location). Hereby we find that the highest number of unique SNPs occurs in samples from Schwäbisch Hall (31 SNPs) dating back to ca 1750 CE. Regarding the total number of SNPs per sampling site we also find that most of the variation identified in the historical material came from the Schwäbisch Hall samples (121 SNP). However, the higher number of variants from this site can be explained with the larger number of samples (five samples). The most distinct samples came from other sites. These were the KP210080 sample from Göttingen and the KP210099 and KP210124 samples from Zaisenhausen/Mulfingen and Weipertshofen, which represented 93, 90 and 80 SNPs, respectively, relative to the rye chloroplast reference genome (**Fig. S1**). The KP210080 sample from Göttingen was collected from the town hall that dates back to the 14th century and it is plausible to speculate that exactly this sample is more distinct because it is older (**Fig. S1)**. However, two other chloroplast sequences that we obtained from the Reutlingen *Wellerhölzer* also date back to the 14th century (KP210196 and KP210197, respectively) and these have a lower number of SNPs compared to all other samples (44 SNPs). Hence, although the low sampling size does not allow conclusive analyses, our analyses show that genetic variation over time has been maintained in rye fields up till the 19^th^ century as illustrated by the variation that we were able to recover in the samples from Zaisenhausen/Mulfingen..

### 3.3. Genetic diversity in *Wellerhölzer* rye is reduced with the intensification of crop cultivation

To gain insights into the evolutionary trajectory of the medieval cereals with specific focus on the medieval rye, we obtained 17 complete chloroplast sequences of historical cereals extracted from the walls of *Wellerhölzer*. We used these sequences to address their phylogenetic relationships with 15 chloroplast sequences of present-day samples (**Fig. 2B**). *Wellerhölzer* samples identified as wheat (KP210077) and barley (KP210078) were grouped with the respective present-day accessions of these two species, thereby confirming the correct archaeobotanical identification. Chloroplast sequences of non-crop *Secale* species and modern rye accessions (Bernhardt et al. 2017; (**Tab. S3**)) clustered together while the historical *Wellerhölzer* chloroplast sequences, including those identified as Cerealia (i.e., cereals not identified to genus or species level) were clustered together. Among the *Wellerhölzer* samples, the phylogenetic analysis clustered samples from the same or near locations together (e.g., *Wellerhölzer* from Schwäbisch Hall and Weipertshofen), thereby reflecting closer genetic relatedness of these samples. However, some samples from the same location did not cluster together suggesting extensive variation in rye populations, not only within, but also between locations. For example, *Wellerhölzer* sample KP210099 from Zaisenhausen/Mulfingen showed higher similarity in the chloroplast sequence to a *Wellerhölzer* sample from Göttingen than to a sample from the geographically much closer Schwäbisch Hall. In summary, the rye/Cerealia *Wellerhölzer* form a separate cluster from the recent rye accessions and the genetic relatedness of these historical samples are in most cases location-dependent.

### 3.4. Genetic variants in historical samples are most abundant in chloroplast genes related to energy metabolism and chloro-respiration

To further investigate the genetic diversity in chloroplast sequences of the historical rye samples, we characterized the distribution of variable sites across the chloroplast genomes and its genetic elements.

Correlating positions of coding sequences and variable positions, we find that genetic variation is not randomly distributed across the chloroplast genomes. Of the total 217 SNPs we find 51 SNPs (23.5%) located in 28 out of the 107 annotated chloroplast genes (26.2%). Of these, 24 SNPs (47%) out of 51 the SNPs in coding sequences resided in genes involved in ATP synthesis and electron transport during photosynthesis that regulate energy metabolism and chloro-respiration 13 SNPs (25.5%) in ATP synthase genes and 11 SNPs (21.5%) in the NADH dehydrogenase-like (ndh) genes, respectively) (**Fig. 3A, 3B**). We further assessed the putative relevance of the called SNPs and annotated their impact as “neutral” or “non-neutral”. We note that of the 217 variable sites, four represent multiple alleles which we defined individually according to their putative impact, i.e. the total number of annotated SNPs was 221. The functional annotations of variants conducted with SnpEff (Cingolani et al., 2012) showed that the identified SNPs were predominantly located in upstream regions of coding sequences including 181 (82%) (**Fig. 3C**). Two SNPs (1%) were located downstream of gene sequences and 22 SNPs (10%) were neutral synonymous variants. Thirteen SNPs (6%) were missense, putatively non-neutral, variants, out of which seven alone were located in the genes encoding the enzymes ATP synthase (atpA, atpB, atpE, atpE, and atpF) and NADH dehydrogenase-like (ndhG, respectively). Additionally, two stop codon mutations (1%) were also present in ATP synthase genes (atpA and atpE), suggesting that the genes involved in cellular metabolism do not only possess higher mutation load in comparison to other chloroplast genes, but harbor higher number of non-neutral polymorphisms.

**Fig. 3.**
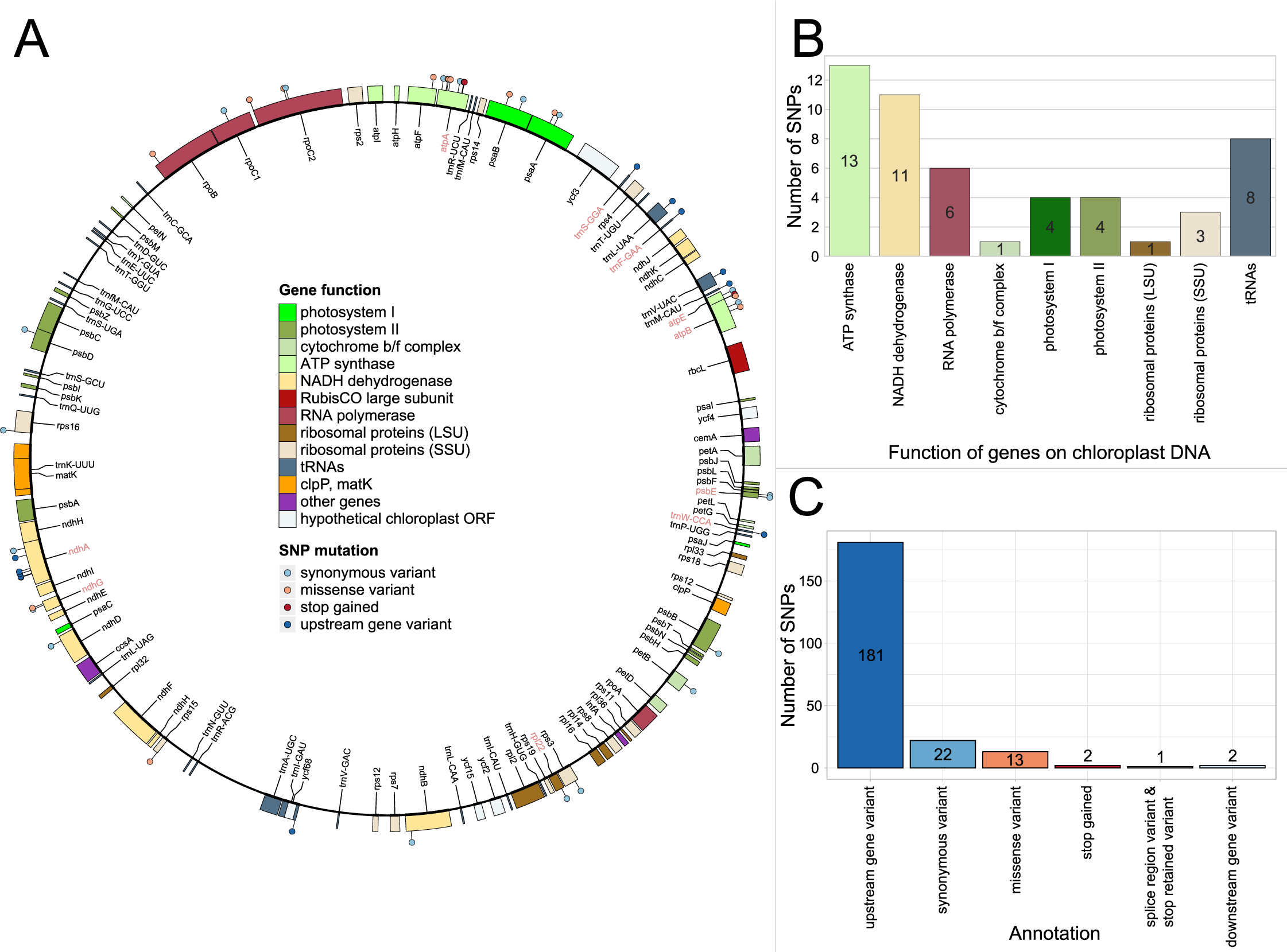
Mutation map in the chloroplast genomes of the historical rye samples. **A)** Distribution of the identified ancient mutations in chloroplast genes from rye varieties sampled from the medieval *Wellerhölzer*. “Lollipop” symbols designate single nucleotide polymorphisms (SNPs) present in respective rye chloroplast genes. Different colors of lollipops designate their functional annotation and gene names. **B)** Bar plot displaying SNP number per 1000 bp window for all chloroplast genes that contain a SNP. The color of the bars designates the chloroplast gene function and corresponds to the legend in A. **C)** Bar plot showing the number of all ancient SNPs in the chloroplast genome (inside and outside chloroplast genes) with their respective functional annotation identified by SnpEff.

Next to recording the total number of SNPs in specific genes, we also calculated the SNP density per gene length (**Fig. 4**). In agreement with our conclusion from crude SNPs numbers, we find that the genes with the highest frequency of variable sites, indeed are genes involved in energy metabolism and chloro-respiration. ATP synthase A (atpA) and NADH dehydrogenase-like A (ndhA) had the highest densities of variable sites (3.5 and 2.3 SNPs per 1000 bp, respectively), while the Rubisco gene and mating type genes (clpP, matk) on the other hand, did not exhibit any variable sites at all. Taken together, our molecular analysis shows the highest variation at the chloroplast DNA level of the historical *Wellerhölzer* rye samples in genes involved in energy production and chloro-respiration, indicating the significant role that these genes might have had in the evolutionary trajectory of rye.

**Fig. 4.**
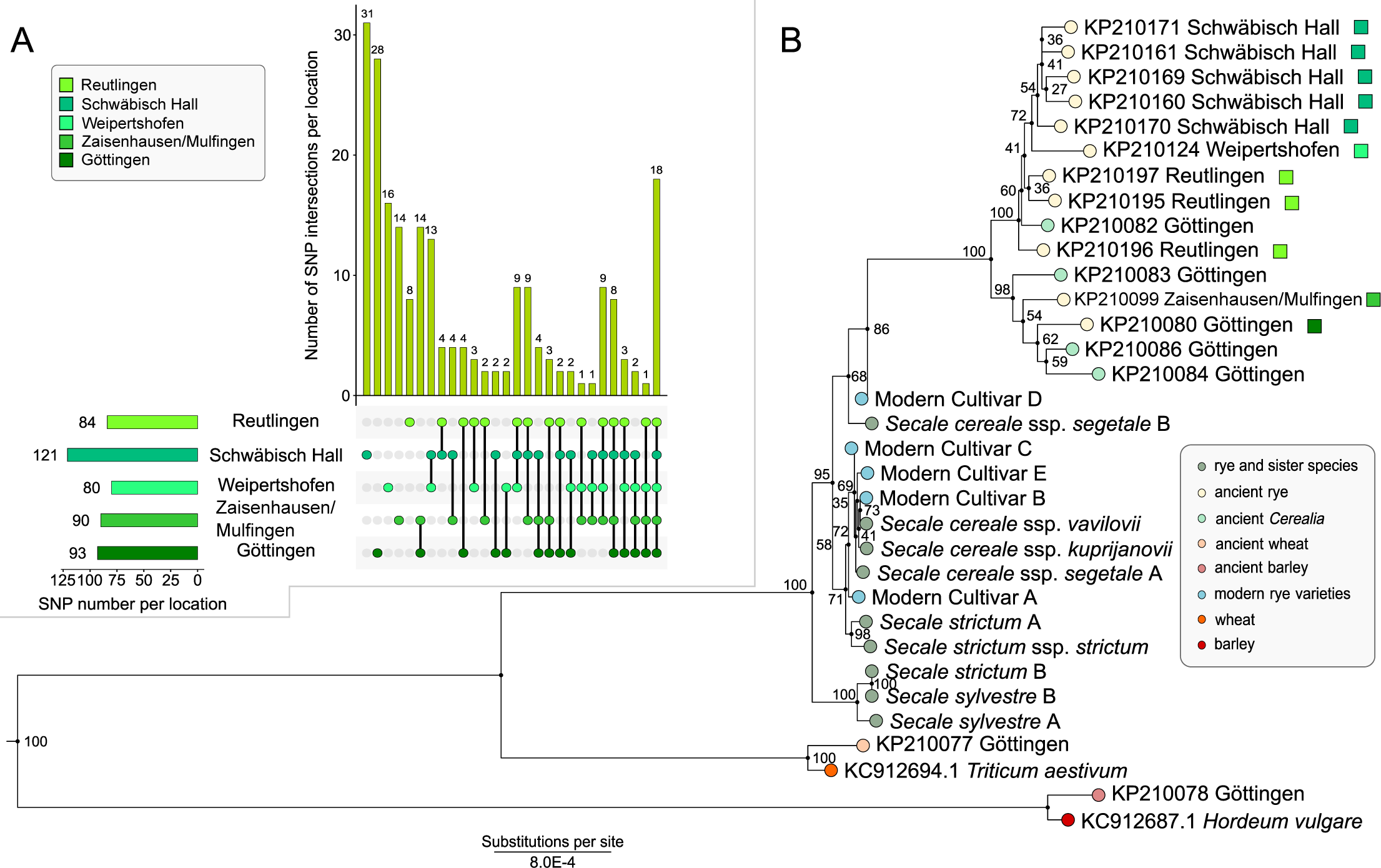
Normalized number of ancient SNPs in rye chloroplast genes. **A)** Number of SNPs per 1000 bp window for each rye chloroplast gene that harbored a mutation. **B)** Barplot displaying the distribution of SNP numbers in ten rye chloroplast genes with the highest SNP load. Different bar colors in A and B correspond to the respective gene function of a gene carrying a mutation.

## 4. Discussion

In our study we made use of unique historical material originating from half-timbered houses at different locations in southern and central Germany to study genetic diversity in historical plant material. The buildings were constructed at different times from the 14^th^ to the 19^th^ centuries, using rye straw and its weedy accompanies for the manufacture of *Wellerhölzer*. The plant remains coiled around the wooden plank were largely incapsulated in the loam covering the *Wellerhölzer* for centuries, and are therefore well-preserved as desiccated, dry matter and thus intact. This allowed us to combine archaeobotanical analyses with ancient DNA sequencing methods to characterize the plant and rye diversity at field sites 200-600 year ago in today’s Germany.

Due to the excellent preservation of the rye straw associated with the *Wellerhölzer*, we were able to sequence the chloroplast genomes of seventeen samples, consisting mainly of samples of rye, but also one sample of barley and one of wheat (**Fig. 2B**). Ancient DNA sequencing of plant material is challenged by the rapid decomposition of plant cell wall structures, which readily occurs under normal environmental conditions. The loam covering the straw has, however, resulted in a desiccated preservation of plant remains, which also sufficiently preserved the plant cell wall structures required for ancient DNA sequencing.

Studies of desiccated plant material from archaeological contexts have previously been conducted successfully on, amongst others, grains of free-threshing wheat (*Triticum aestivum* and *T. durum*), broomcorn millet (*Panicum miliaceum*), barley (*Hordeum vulgare*) and sorghum (*Sorghum bicolor)* (Li et al., 2016; Mascher et al., 2016; Oliveira et al., 2012; Smith et al., 2019). In the case of barley, 6000 year old specimens were preserved inside a cave in the Judean desert aided by the hot and arid conditions of the surroundings (Mascher et al., 2016). For this ancient barley material, exome capture data from six grains provided unique insights into the genetic composition of barley that was cultivated by Neolithic farmers. Intriguingly, comparing genetic information from a large number of modern barley varieties showed that already in the Neolithic, domesticated barley was considerably distinct from wild barley. This suggests a significant genetic modification already from the onset of barley domestication in the Fertile Crescent.

In contrast to the 6000-year-old barley remains, we find a significant variation in the chloroplast genome of the historical rye samples from today’s Germany. We mapped sequencing reads to the reference chloroplast genome of a modern rye variety *S. cereale* cv. Imperial (Middleton et al., 2014), and identified 217 variable sites among the historical rye samples. Notably, the samples from Schwäbisch Hall showed a large number of unique variable sites in the chloroplast genome, although these samples are among the more “modern” samples (ca year 1750). In general, most of genetic variants that we identify in the historical samples are unique and not present among the modern varieties that we include for a comparison. This observation most likely reflects the intensification of rye breeding during modern times and consequently the loss of past genetic variation. From the Medieval period onwards (ca. 8^th^ century CE) and throughout the 19^th^ century, the three-field system became a common cultivation practice within the modern borders of Germany (Miedaner, 2014). Hereby, one part is sown with a summer crop, another part is sown with a winter crop, the final part is left fallow, and each part is rotated every season (Jones, 2007; Miedaner, 2014). Using the mouldboard plough, wet autumn soils were turned, enabling the cultivation of winter crops such as rye (Jones, 2007; Miedaner, 2014). Until the 1850’s, local selection processes resulted in a diverse set of varieties of rye, while from the 1850”s onward, selection processes targeted at specific characteristics of rye became more generalized, for example leading to the first variety of rye that was more resistant to lodging (Schlegel, 2014). Our archaeobotanical analysis suggests that the rye from all investigated locations were likely cultivated as part of a three-field system. Based on the high genetic diversity of the samples included in our investigation, we can speculate that the samples represent varieties of rye that were the result of local selection processes, rather than targeted breeding.

We used the sequence data to investigate the distribution of historical genetic variants in relation to the chloroplast gene annotation. Our results show an increased number of genetic variants in the historical samples in the chloroplast genes that code for subunits of large protein complexes involved in energy production and chloro-respiration, primarily the ATP synthase and the NADH dehydrogenase-like (ndh) protein complex. The chloroplast ATP synthase produces the ATP needed for photosynthesis and plant growth (Yamamoto et al., 2023), and the distinct haplotypes observed in the ATP synthase genes of the *Wellerhölzer* rye potentially reflects later breeding efforts to increase crop biomass (Galán et al., 2021). Furthermore, the chloroplastic ATP synthase complex also plays a role in plant stress response, since polymorphisms in the subunits of this complex have been associated with cold stress tolerances in cucumber (Oravec and Havey, 2021). The chloroplast gene with the highest number of genetic variants, atpA, was found to be associated with increased resistance to the fungal pathogen *Botrytis cinerea* in tomato (Gong et al., 2021). ATP synthase genes is in general considered to play a significant role in plant stress tolerance and may therefore also have evolved in response to distinct environmental conditions and stressors. Chloroplast genes coding for subunits of the NDH complex involved in cyclic electron transport for photosystem I (PSI) and chlororespiration, also exhibited an increased number of genetic variants in the historical rye samples. The role of the NDH subunit encoding genes in abiotic stress alleviation has been demonstrated in several important crop species, such as tobacco and rice, where specific mutations in different NDH subunits were associated with improved response to heat and cold stress (Wang et al., 2016), as well as to attenuation of oxidative stress under fluctuating light conditions (Yamori et al., 2011). Taken together, the highest portion of the identified point mutations in the chloroplasts of historical *Wellerhölzer* rye locate in genes involved in abiotic and biotic stress responses, and the distribution of these mutations in ancient chloroplast genomes possibly reflects non-targeted footprints of rye domestication and breeding.

In summary, our study demonstrates the great potential of combining aDNA sequencing and archaeobotanical analyses to address the breeding and cultivation history of crops. We here describe the plant diversity in rye fields at five rural locations in Germany. Our sampling include specimen from the 14^th^ to the 19^th^ century and illustrates a considerably change in biodiversity and genetic composition of rye varieties. In general, we observe a considerable loss of genetic diversity in present day rye cultivars compared to historical rye varieties. This observation is in line with the development of agricultural practices and breeding efforts. Present day and future crop production is challenged by the consequences of climate change, including drought stress, flooding and altered spectra of pathogens. Breeding efforts should aim to restore and preserve genetic variation in crops to increase resilience and adaptability of plant populations. In this regard, insights into the past spectrum of genetic diversity in cereal crops may point to ancient stress resistances that can be reestablished in present day crops to meet future challenges.

## Supporting information

KomluskiSupplementary

## Acknowledgements

This work was funded through the DFG-funded Excellence Cluster ROOTS (EXC2150). We are grateful for support by PD Dr. Elena Marinova and Dr. Elske Fischer from the Archaeobotanical Laboratory of the State Office for Cultural Heritage Baden Württemberg, Hemmenhofen/Gaienhofen, Germany.

## 5. Author Contributions

**JK**-Conceptualization; Formal analysis; Visualization; Writing – original draft.

**EHS** - Conceptualization, Writing and editing of the manuscript, Project management

**SF** - Conceptualization, Writing and editing of the manuscript

**WK -** Conceptualization, Writing and editing of the manuscript

**FS -** Writing and editing of the manuscript

**BKK** – Conceptualization, data generation, editing of the manuscript

**BC** – Data generation

**MF** – Data generation, editing of the manuscript

## Supplementary Materials and Appendices

**Fig. S1.** Intersections between ancient SNPs isolated from the historical rye samples. Upset plot showing shared ancient SNP between different rye samples from different sampling locations. Different shades of green correspond to sampling location.

**Table S1.** List of ancient cereal varieties with their 14C dating used in this study.

**Table S2.** Overview of autecological characteristics and functional attributes collected for the taxa of arable weeds and wild plants from the Wellerholz from Schwäbisch Hall. The information was gathered from Ellenberg (1988) and Biolflor (Klotz et al. 2002).

**Table S3.** List of chloroplast genomes of modern cereal varieties and rye sister species that were used for phylogenetic analysis.

**Appendix 1** Description of archaeobotanical analyses.

**Appendix 2** Scripts used for sequence analyses

