## Supplementary material for "Sequencing of historical plastid genomes reveal exceptional genetic diversity in early domesticated rye plants": KomluskiSupplementary: Appendix_1.docx

**Title:** Sequencing of historical plastid genomes reveal exceptional genetic diversity in rye before the start of breeding

Jovan Komluski, Sofia Filatova, Frank Schlütz, Benjamin Claaßen, Manfred Rösch, Ben Krause-Kyora, Wiebke Kirleis, Eva H. Stukenbrock

**Appendix 1: Archaeobotanical analyses**

**Methods**

For the analysis of archaeobotanical remains obtained from the Wellerholz from Göttingen, including sample processing sorting and quantification, we used our previously established pipeline for the Wellerhölzer material (Filatova et al., 2021). The details on the analysis of archaeobotanical remains obtained from the Wellerholz from Schwäbisch Hall conducted by Manfred Rösch and Elske Fischer at the archaeobotanical department of the State Office for Cultural Heritage Baden Württemberg in Hemmenhofen/Gaienhofen (Germany) are published in Rösch and Fischer (1997).

Cultivation practices and soil conditions of the rye harvests represented in the two *Wellerhölzer* were reconstructed based on a combination of data on 1) the autecology and 2) the functional autecology of the identified arable weed taxa (for a more elaborate discussion, see Filatova et al. (2021)). These two approaches are based on the notion that characteristics of plants are indicative of certain climatic conditions (e.g., temperature and light), soil conditions (e.g., pH and nitrogen), and activities of soil disturbance (e.g., digging and trampling) (Bogaard et al., 1999; Charles et al., 2002; Duckworth et al., 2000; Ellenberg, 1988). (A Bogaard et al., 1999; M Charles et al., 1997; Mike Charles et al., 2002; Duckworth et al., 2000; Ellenberg, 1988). Such data can be used to indicate a range of plant-cultivation practices and soil conditions, such as sowing time, the practice of weeding, and the use of fertilizer. Synecological information of the arable weeds and wild plants in the samples was used to indicate the contemporary plant sociological groups of the taxa.

All taxa that were identified to species level were included in the analysis following a qualitative approach. Functional attributes and autecological characteristics of the selected species were used to reconstruct cultivation practices (Table A.1). Data on synecology, autecology, and functional attributes were collected from Ellenberg (1988) and Kühn et al. (2004)(Klotz et al., 2002).

**Table A.1** Autecological characteristics and functional attributes used to indicate cultivation practice. The criteria used to define flowering onset and duration are: Early flowering onset: January-April; Intermediate flowering onset: April-June; Late flowering onset: July or later; Short flowering duration: 1-3 months; Medium flowering duration: 4-5 months; Long flowering duration: > 5 months. The table and flowering onset/duration criteria are adapted from Hillman 1981, Kreuz and Schäfer 2011, Neveu et al. 2021, Charles et al. 2002, Jones et al. 2005, Bogaard et al. 2005, Van der Veen 1992 and Leuschner and Ellenberg 2017.

| **Cultivation practice** | | **Associated (aut)ecological characteristic** |
| --- | --- | --- |
| Sowing season | Autumn sowing | Early flowering onset |
|  |  | Intermediate flowering onset |
|  |  | Short flowering duration |
|  |  | Long flowering duration |
|  | Spring sowing | Late flowering onset |
|  | Inconclusive | Medium flowering duration |
| Disturbance degree | High disturbance | Long flowering duration |
|  |  | Ruderal strategy type |
|  |  | Therophyte lifeform |
|  |  | Annual with reproduction by seed |
|  |  | Perennial with subterranean reproductive organs |
|  | Low disturbance | Perennial without subterranean reproductive organs |
|  |  | Short flowering duration |
| Ploughing season | Autumn ploughing | Early flowering onset and/or short flowering duration |
|  | Spring ploughing | Late flowering onset |

**Results**

The material from the *Wellerholz* from Schwäbisch Hall (dating to 1750 AD) comprised 829 plant remains. The majority were remains of cereals (51%; mainly rye ears) followed by arable weeds and wild plants (44%), the latter of which included 16 taxa that were identified to species level (Table A.2 and Table S3). Most of these species thrive in the modern segetal plant communities of the Chenopodietea (3.3 in Table S3) and the Secalietea (3.4 in Table S3) as decribed in Ellenberg (1988). The dominant lifeform is therophyte-hemicryptophyte (n=10; Fig. A.1), followed by therophyte (n=3), hemicryptophye (n=1) and hemicryptophyte-geophyte (n=1). Twelve species have an annual life lifecycle, three a perennial one, and one species can thrive both as an annual as well as a perennial. An intermediate (n=12) flowering onset is most common, followed by early (n=3) and late (n=1), while a short flowering duration prevails (n=7), followed by long (n=5) and medium (n=4). Species that have both a competitive and a ruderal ecological strategy as well as species that solely have a ruderal strategy are equally represented (n=6), and two different combinations of competitors, ruderals and stress-tolerators occur in one occasion each.

**Table A.2** Overview of plant taxa identified to the level of species from the Wellerholz from Schwäbisch Hall (Rösch and Fischer, 1997).

| **Taxon original** | **Absolute find quantity** |
| --- | --- |
| *Agrostemma githago* | 1 |
| *Alopecurus myosuroides* | 17 |
| *Apena spica-venti* | 33 |
| *Chenopodium album* | 3 |
| *Fragaria vesca* | 1 |
| *Matricaria chemomilla* | 1 |
| *Myosodon aquaticum* | 2 |
| *Odontites vernus* | 3 |
| *Papaer rhoeas* | 1 |
| *Poa annua* | 2 |
| *Polygonum aviculare* | 258 |
| *Rubus idaeus* | 1 |
| *Senecio sylvaticus* | 1 |
| *Senacio vulgaris* | 2 |
| *Sonchus oleraceus* | 13 |
| *Stellaria media* | 1 |

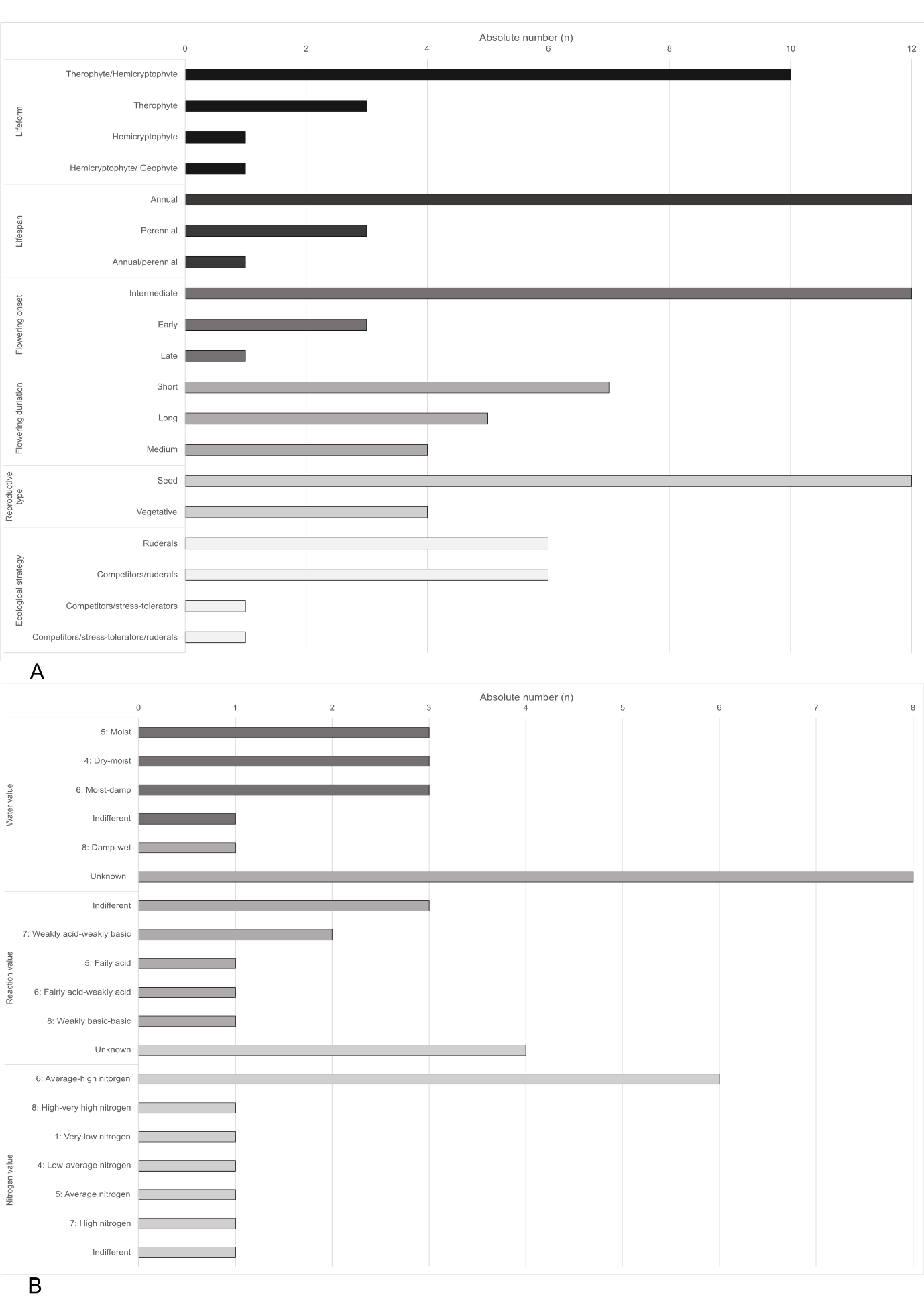

**Fig. A.1** Bar chart summarizing the autecological data and functional attributes of the arable weeds and wild plants from Schwäbisch Hall that were used to characterize the sowing time and cultivation intensity of the rye (the data collected from Ellenberg (1988) and Biolflor (Klotz et al. 2002). **b.** Bar chart summarizing the edaphic factors used to characterise conditions of the soil based on the arable weeds and wild plants from Schwäbisch Hall (values collected from Ellenberg 1988).

According to the criteria summarized in Table A.1, an early and intermediate flowering onset with a short and long flowering duration are typical indicators of autumn sowing. The prevalence of therophyte lifeforms with an annual reproduction cycle and a ruderal strategy type indicate a high level of disturbance, and the early flowering onset and short flowering duration imply autumn ploughing. The co-occurrence of nearly equal amounts of short and long flowering durations renders these two characteristics inconclusive with respect to the definition of the degree of disturbance. Taken together, these data strongly suggest that the rye was sown in autumn, and thus grew as a winter rye, and that the arable weeds and wild plants were adapted to a high degree of disturbance.

Ellenberg’s indicator values of edaphic factors show a diverse range of values related to the arable weeds and wild plants from Schwäbisch Hall (Fig. A.b). The water value is dominated by moist indicators (n=5), followed by dry-moist (n=3), and moist-damp (n=3) indicators. One species thrives in damp-wet soils, while three are indifferent and one is unknown. The majority of the species are indifferent to soil pH (n=8), three are indicative of weakly acidic-weakly basic conditions, two of fairly acidic conditions, while fairly acidic-weakly acidic and weakly basic conditions are indicated by one species each. High to very high nitrogen content is indicated by six species, average to high nitrogen content is indicated by four species, and all other values (ranging from very low to high nitrogen content) are indicated by one species each. In summary, the indicator values suggest moist soil conditions, a high nitrogen content and more acidic than basic conditions. Reaction values show intermediate to acid conditions since many species are indifferent. Many indicate basic soils and species with indication for acid soils can grow on basic soils as well. The high representation of species growing on acid soils however may hint to originally poor soils with low pH that were manured. It has been proposed that, at least from the medieval period onward, rye was typically grown on more acidic soils than species of wheat, which corresponds well to the conditions reflected by the indicator values (Rösch and Fischer, 1999). The relatively high nitrogen content might be indicative for manuring of the field. Furthermore, the diversity of indicator values represented by the species of arable weeds and wild plants suggest a low level of competition in the field, which is characteristics for ruderals thriving in highly disturbed habitats (Grime, 1977).

In summary, the reconstruction of cultivation practices and soil conditions of the material from Schwäbisch Hall indicates that the rye was sown as a winter cereal in highly disturbed fields with a moist, acidic to neutral soil that was likely manured. These conditions are similar to the ones reflected by the material from Göttingen, with the exception of the nitrogen content of the soil, which in the case of Göttingen did not suggest the practice of manuring.

Van Der Veen, M. Crop Husbandry Regimes: An Archaeobotanical Study of Farming in Northern England; Sheffield Archaeological Monographs: Sheffield, UK, 1992.
