## Supplementary material for "Sequencing of historical plastid genomes reveal exceptional genetic diversity in early domesticated rye plants": KomluskiSupplementary: Appendix_2.pdf

```
Paper title: Sequencing of historical plastid genomes reveal exceptional genetic diversity in
             rye before the start of breeding

Code author: Jovan Komluski

Date: 15/03/2024
Appendix: The following document contains six scripts that have been used for variant calling,
         variant annotation, molecular analysis and data visualization of the medieval rye
         DNA dataset:
          01_variant_calling.sh
          02_variant_annotation.py
          03_SNP_per_gene.sh
          04_nucleotide_diversity.py
          05_generating_upset_and_circos_plotting_dataframes.py
          06_upset_and_circos_plots.R
```

```
---------------------------------01_variant_calling.sh---------------------------------------

Purpose: bash script for calling SNPs from the reads mapped to a chloroplast genome
         The output of the script is a single merged vcf file with variants from all
         studied isolates.
         As an input script uses text file with the paths of the bam file locations
         to call SNPs from the corresponding bam files that were generated via EAGER
         pipeline (https://doi.org/10.1186/s13059-016-0918-z).

Example of the first line of the input text file with the list of bam files for variant
    calling:
/path/to/bam/file/KP210080_21Jun170_D06_S42_L001/KP210080_21Jun170_D06_S42_L001_rmdup.bam

Software prerequisites:
samtools 1.7
bcftools 1.7
tabix (htslib) 1.10.2
```

```
#!/bin/bash

setting up paths and variables
INPUT_BAM=/path/to/list_of_bam_files_that_passed_quality_control_and_duplicates_removal.txt
REF_FASTA=/path/to/reference/chloroplast/genome.fasta
OUTDIR_RMDUP=/path/to/output/directory/

function for iterating over bam files of interest
snp_calling () {
for FILE in $(cat ${INPUT_BAM});

do

```

```
FILE1=${FILE%_rmdup.bam}

FILE_BASE=$(basename ${FILE%_rmdup.bam})


sorting of bam files
samtools sort -@ 10 ${FILE} -o ${FILE1}.sorted.bam

mapping of the sorted bam files to the reference genome
samtools mpileup -C 50 -Q 0 -q 20 -uf \
${REF_FASTA} \
${FILE1}.sorted.bam | \
bcftools call -vc -O u -o ${FILE1}.bcf

conversion of bcf to vcf file and filtering for biallelic SNPs with mapping quality higher
    than 20
bcftools view -i 'QUAL>20' \
--exclude-types indels \
--max-alleles 2 ${FILE1}.bcf \
-o ${OUTDIR_RMDUP}${FILE_BASE}.vcf

compressing of the vcf file
bgzip ${OUTDIR_RMDUP}${FILE_BASE}.vcf

indexing of the compressed vcf file
tabix -p vcf ${OUTDIR_RMDUP}${FILE_BASE}.vcf.gz

done

}

snp_calling
```

```
merging of all generated vcf files into a single vcf file
bcftools merge -o ${OUTDIR_RMDUP}merged_isolates.vcf -O z *.vcf.gz
-----------------------------------------------------------------------------------------------
```

```
----------------------------------02_variant_annotation.py--------------------------------------

Purpose: Extracting SNP annotation information from a ann.vcf file
         produced by SnpEff and generating plotting dataframe that shows
         proportion of different annotations among SNPs identified in
         chloroplast genome of historical rye isolates
```

```
importing necessary modules
from os import sep
import pandas as pd
```

```
import numpy as np


setting input file directory path
vcf_path = '/path/to/merged/vcf/file/with/SNPs/from/all/isolates/'

removing all of the lines before the line of a vcf file that starts with #CHROM
with open(vcf_path + 'merged_vcf_file.vcf') as infile:
    for n,row in enumerate(infile):
        if row.startswith("#CHROM"):
            break

importing the merged vcf file with SNPs with #CHROM line as header
df_snps = pd.read_csv(vcf_path + 'merged_vcf_file.vcf', sep="\t", skiprows=n)

the code below extracts the first annotation from the INFO column for each SNP
df_snps = df_snps.iloc[:, 0:9]
df_snps[['INFO','ANN_SNPEFF']] = df_snps['INFO'].str.split('ANN=', n=2, expand=True)
df_snps['ANN_SNPEFF'] = "," + df_snps['ANN_SNPEFF'].astype(str)
df_snps['A_ANN'] = df_snps['ANN_SNPEFF'].str.split(r',A\|').str[1].str.split(r'\|').str[0]
df_snps['C_ANN'] = df_snps['ANN_SNPEFF'].str.split(r',C\|').str[1].str.split(r'\|').str[0]
df_snps['T_ANN'] = df_snps['ANN_SNPEFF'].str.split(r',T\|').str[1].str.split(r'\|').str[0]
df_snps['G_ANN'] = df_snps['ANN_SNPEFF'].str.split(r',G\|').str[1].str.split(r'\|').str[0]

df_snps.fillna('!', inplace=True)

df_snps['annotation'] = df_snps['A_ANN'].astype(str) + ";" +
                        df_snps['C_ANN'].astype(str) + ";" +
                        df_snps['T_ANN'].astype(str) + ";" +
                        df_snps['G_ANN'].astype(str) + ";"
df_snps['annotation'] = df_snps['annotation'].str.strip().str.replace('!;','')

df_snps.drop(['ANN_SNPEFF', 'A_ANN', 'C_ANN', 'T_ANN', 'G_ANN'], axis=1, inplace=True)


df_snps['annotation'] = df_snps['annotation'].str.split(";").map(set).str.join("|")
df_snps['annotation'] = df_snps['annotation'].str.replace('|','', n=1)

making a dataframe in which SNP annotations will be counted per annotation category
df_counting = df_snps.assign(type =
    df_snps['annotation'].str.split(r'\|')).explode('annotation')

preparing the dataframe for circos plot of chloroplast genome-wide SNP distribution
df_snps['color'] = 'NaN'
df_snps.loc[df_snps['annotation'] == 'downstream_gene_variant', 'color'] = '#D1E5F0'
df_snps.loc[df_snps['annotation'] == 'missense_variant', 'color'] = '#F4A582'
```

```
df_snps.loc[df_snps['annotation'] == 'splice_region_variant&stop_retained_variant', 'color'] = '#FDDBC7'
df_snps.loc[df_snps['annotation'] == 'stop_gained', 'color'] = '#B2182B'
df_snps.loc[df_snps['annotation'] == 'synonymous_variant', 'color'] = '#92C5DE'
df_snps.loc[df_snps['annotation'] == 'upstream_gene_variant', 'color'] = '#2166AC'
df_snps.loc[df_snps['color'] == 'NaN', 'color'] = '#ED9ED6'

df_snps["end"] = df_snps["POS"] + 1
df_snps["start"] = df_snps["POS"]

df_snps = df_snps.rename(columns={'#CHROM': 'chr'})
df_snps = df_snps.rename(columns={'POS': 'position'})
df_snps = df_snps.rename(columns={'annotation': 'type'})

df_snps = df_snps[['chr', 'position', 'start', 'end', 'type', 'color']]

exporting the dataframe with SNP coordinates for circos plot
df_snps.to_csv(vcf_path + 'circos_snps.csv', sep='\t', index=False)

generating empty lists where specific SnpEff annotation to store annotation type and count
ann_list = []
count_list = []

def counting_number_of_annotations():
Function that stores annotations to list, counts number of respective annotation and stores it to a list
    for annotation in df_counting['annotation'].unique().tolist():
        ann_list.append(annotation)
        count_list.append(df_counting['annotation'].value_counts()[annotation])

counting_number_of_annotations()
```

```
creating the dataframe with number of SnpEff annotations for plotting (Fig. 3C)
df_annotcount = pd.DataFrame({'variant_annotation': ann_list, 'nb_snp': count_list,'color': ''})
df_annotcount['nb_snp'].sum()

df_annotcount.loc[df_annotcount['variant_annotation'] == 'downstream_gene_variant',
                                                                  'color'] = '#D1E5F0'
df_annotcount.loc[df_annotcount['variant_annotation'] == 'missense_variant',
                                                                  'color'] = '#F4A582'
df_annotcount.loc[df_annotcount['variant_annotation'] == 'splice_region_variant&stop_retained_variant',
                                                                  'color'] = '#FDDBC7'
df_annotcount.loc[df_annotcount['variant_annotation'] == 'stop_gained',
                                                                  'color'] = '#B2182B'
```

```
df_annotcount.loc[df_annotcount['variant_annotation'] == 'synonymous_variant',
                                                                   'color'] = '#92C5DE'
df_annotcount.loc[df_annotcount['variant_annotation'] == 'upstream_gene_variant',
                                                                   'color'] = '#2166AC'

export count dataframe for plotting
df_annotcount.to_csv(vcf_path + 'annotation_count_df.csv', sep='\t', index=False)
-----------------------------------------------------------------------------------------------
```

```
----------------------------------03_SNP_per_gene.sh------------------------------------------

Purpose: bash script that takes the gff3 file of the reference chloroplast genome
         in [NCBI format] and a vcf file produced by user as an input to:
         1) identify number of SNPs in the respective chloroplast genes with the mutation
         2) divide chloroplast genes into 1000 bp windows
         3) calculate the SNP number per window for each of the chloroplast genes
         4) makes an input for script that generates file for circos plotting dataframe

         Software prerequisites: bedtools v2.26.0
```

```
#!/bin/bash

setting up paths and input files
READDIR=/path/to/input/gff3&vcf/files/
OUTDIR=/path/to/script/output/directory/

ENTRY_GFF=reference_chloroplast_genome_annotation.gff3
ENTRY_VCF=merged_vcf_file_with_variants_from_all_isolates.vcf


converting gff3 file of the reference chloroplast genome to the bed file with the chloroplast gene coordinates
awk '!/^#/ {if ($3=="gene") {print "KC912691" "\t" $4 "\t" $5 "\t" $9} }' ${READDIR}${ENTRY_GFF} > ${OUTDIR}KC912691_chloroplast.bed


making a bed file out of the merged vcf.gz file with mutations from all of the isolates
zless ${READDIR}${ENTRY_VCF} | awk '!/^#/ {print $1 "\t" $2 "\t" $2+1}' > ${OUTDIR}ancient_unique_snps.bed

intersecting of the bed file with the SNP coordinates from the ancient isolates with bed file with gene coordinates from the reference chloroplast genome
bedtools intersect -a ${OUTDIR}ancient_unique_snps.bed -b ${OUTDIR}KC912691_chloroplast.bed -wb > ${OUTDIR}aSNPs_in_genes.bed

calculating the chloroplast gene sizes and removing the duplicates
```

```
awk '{print $0 "\t" $6-$5}' ${OUTDIR}aSNPs_in_genes.bed | awk '!seen[$2,$3,$5,$6,$7]++' > ${OUTDIR}aSNPs_in_genes_with_gene_sizes.bed

generating the list of SNPs with an ancient SNP
awk '{print $7}' ${OUTDIR}aSNPs_in_genes.bed | sort | uniq | \
awk -F';' '{print $2}' | awk -F'=' '{print $2}' > ${OUTDIR}list_of_genes_with_aSNP.csv

generating a dataframe with three columns (annotation column of a gene, SNP number per gene; gene name)

generating_df () {
for f in $(awk '{print $7}' ${OUTDIR}aSNPs_in_genes_with_gene_sizes.bed | sort | uniq);

do

echo $f >> ${OUTDIR}nb_aSNP_per_gene_list.csv;
awk -v var="$f" '{ if ( $7 == var ) { print $0; } }' ${OUTDIR}aSNPs_in_genes_with_gene_sizes.bed | wc -l >> ${OUTDIR}nb_aSNP_per_gene_list.csv

done

}

generating_df

awk '{printf "%s%s",$0,NR%2?"\t":RS}' ${OUTDIR}nb_aSNP_per_gene_list.csv > ${OUTDIR}tmp \
&& mv ${OUTDIR}tmp ${OUTDIR}nb_aSNP_per_gene_list.csv

paste ${OUTDIR}nb_aSNP_per_gene_list.csv ${OUTDIR}list_of_genes_with_aSNP.csv > ${OUTDIR}nb_of_aSNP_per_gene.csv

inserting column names to a csv file with number of SNPs per chloroplast gene
sed  -i '1i attribute\tnb_aSNP\tgene' ${OUTDIR}nb_of_aSNP_per_gene.csv

separating each gene in the chloroplast genome into 1000 bp windows
bedtools makewindows -b ${OUTDIR}KC912691_chloroplast.bed -w 1000 -i src > ${OUTDIR}KC912691_chloroplast_allgenes.windows.bed

intersecting 1000 bp gene windows with snp coordinates of snps in genes
bedtools intersect -a ${OUTDIR}aSNPs_in_genes.bed -b ${OUTDIR}KC912691_chloroplast_allgenes.windows.bed -wb > ${OUTDIR}snps_per_gene_window.bed

getting the number of windows per gene
calculating_windows_nb_per_gene () {
for f in $( awk '{print $4}' ${OUTDIR}KC912691_chloroplast.bed);
do
```

```

echo $f >> ${OUTDIR}number_of_windows_per_gene.csv;

awk -v var="$f" '{ if ($4==var) {print $0;} }' ${OUTDIR}KC912691_chloroplast_allgenes.windows.bed | wc -l >> ${OUTDIR}number_of_windows_per_gene.csv;

done

}

calculating_windows_nb_per_gene

awk '{printf "%s%s",$0,NR%2?"\t":RS}' ${OUTDIR}number_of_windows_per_gene.csv > ${OUTDIR}tmp && mv ${OUTDIR}tmp ${OUTDIR}number_of_windows_per_gene.csv

generating file with the chloroplast gene coordinates that will be input for 05_generating_upset_and_circos_plotting_dataframes.py
awk '!/^#/ {if ($3=="gene") {print $1 "\t" $4 "\t" $5 "\t" $9} }' ${READDIR}${ENTRY_GFF} | awk '{sub(/;gene_biotype.*/, "", $4)} 1' | awk '{sub(/.*gene=/, "", $4)} 1' | awk '{print $1 "\t" $2 "\t" $3 "\t" $4}' > ${OUTDIR}KC912691_genes.bed
----------------------------------------------------------------------------------------------
```

```
-----------------------------------04_nucleotide_diversity.py----------------------------------

Purpose: Python script that takes dataframes (df1; df2) from 03_SNP_per_gene.sh script as input to perform:
         normalization of SNP number per 1000 bp gene window;
         calculation of SNP number per gene functional group;
         plotting of Fig. 3B; Fig. 4A; Fig. 4B.

         Input dataframe df1 consists of three columns ("attribute", "nb_aSNP", and "gene" column).
         The first column of the df1 file is the column number 9 from the reference annotation file
         (i.e., KC912691.gff3) for each chloroplast gene.
         The second column represents number of SNPs per respective gene.
         The third column is shortened name of each respective gene to the first 3 characters
         (e.g., for "atpA" gene the value of the column 3 will be "atp").
         The third column allows grouping of SNPs per gene functional group.

         Input dataframe df2 consists of 8 columns:
         column 1 and 4 are reference genome names;
         column2 and 3 are start and end SNP position;
         column 5 and 6 are start and end gene position;
         column 7 is the annotation column from the reference annotation file KC912691.gff3;
```

```
        column 8 is gene size.
```

```
importing necessary modules
from operator import index
from os import sep
from re import T
from turtle import color
from urllib.parse import ParseResultBytes
import pandas as pd
import numpy as np
import seaborn as sns
import matplotlib.pyplot as plt
import matplotlib as mpl
from  matplotlib.ticker import FuncFormatter
from matplotlib import rcParams

setting input file directory path
path = '/path/to/files/that/are/output/of/03_SNP_per_gene.sh/'

importing dataframes
df1 = pd.read_csv(path + 'nb_of_aSNP_per_gene.csv', sep = "\t")
df2 = pd.read_csv(path + 'aSNPs_in_genes_with_gene_sizes.bed', sep = "\t", header=None)

adding column names since input df2 has none
df2.columns =
    ['ref','snp_start','snp_end','REF','gene_start','gene_end','attribute','gene_size']
removing columns with the reference chloroplast genome names
df2.drop(columns=['REF','ref'], inplace=True)

merging df1 and df2 for lines that have the same value in the attribute column
df3 = pd.merge(df1, df2, on=['attribute'], how="outer", indicator=True)
removing duplicated values from the attribute column in the merged dataframe (df3)
df4 = df3.drop_duplicates(subset=['attribute'], keep='first')
removing the indicator column from the dataframe generated in the step above (df4)
df5 = df4.iloc[: , :-1]
moving the "gene_size" column to the position 3 in df5
column_to_move = df5.pop("gene_size")
df5.insert(3, "gene_size", column_to_move)

adding the gene function column to df5
df5['gene_function'] = df5['gene'].str[:3]
adding gene function for the trnk-UUU gene since it belongs to mat gene family
df5.loc[df5['gene'] == 'trnK-UUU', 'gene_function'] = 'mat'
adding gene function for the photosystem I (PSI) genes
df5.loc[(df5['gene'] == 'psaA') |
        (df5['gene'] == 'psaB') |
```

```
        (df5['gene'] == 'psaC') |
        (df5['gene'] == 'psaJ') |
        (df5['gene'] == 'psaL') |
        (df5['gene'] == 'psaF') |
        (df5['gene'] == 'psaE') , 'gene_function'] = 'psaI'

calculating SNP number for genes that are belong to the same gene family
grouped_functions = df5.groupby(by='gene_function')['nb_aSNP'].sum()
grouped_functions_df = grouped_functions.reset_index()

adding annotation column for each gene functional group (family)
grouped_functions_df['families'] = ''
grouped_functions_df.loc[grouped_functions_df['gene_function'] == 'atp', 'families'] = 'ATP
    synthase'
grouped_functions_df.loc[grouped_functions_df['gene_function'] == 'ycf', 'families'] =
    'hypothetical chloroplast ORF'
grouped_functions_df.loc[grouped_functions_df['gene_function'] == 'ndh', 'families'] = 'NADH
    dehydrogenase'
grouped_functions_df.loc[grouped_functions_df['gene_function'] == 'pet', 'families'] =
    'cytochrome b/f complex'
grouped_functions_df.loc[grouped_functions_df['gene_function'] == 'psb', 'families'] =
    'photosystem II'
grouped_functions_df.loc[grouped_functions_df['gene_function'] == 'rpl', 'families'] =
    'ribosomal proteins (LSU)'
grouped_functions_df.loc[grouped_functions_df['gene_function'] == 'rps', 'families'] =
    'ribosomal proteins (SSU)'
grouped_functions_df.loc[grouped_functions_df['gene_function'] == 'rpo', 'families'] = 'RNA
    polymerase'
grouped_functions_df.loc[grouped_functions_df['gene_function'] == 'trn', 'families'] = 'tRNAs'
grouped_functions_df.loc[grouped_functions_df['gene_function'] == 'psaI', 'families'] =
    'photosystem I'
grouped_functions_df.loc[grouped_functions_df['gene_function'] == 'psa', 'families'] =
    'photosystem II'
grouped_functions_df.loc[grouped_functions_df['gene_function'] == 'rbc', 'families'] =
    'RubisCO large subunit'
grouped_functions_df.loc[(grouped_functions_df['gene_function'] == 'ccs') |
                         (grouped_functions_df['gene_function'] == 'cem') |
                         (grouped_functions_df['gene_function'] == 'inf'), 'families'] =
                             'other genes'
grouped_functions_df.loc[(grouped_functions_df['gene_function'] == 'mat') |
                         (grouped_functions_df['gene_function'] == 'clp'), 'families'] =
                             'clpP, matK'
```

```
-----------------------normalization of SNP number per gene per 1000 bp length----------------

         Input dataframes df_snpwin and df_nbwin are outputs from the 03_SNP_per_gene.sh
```

```
        script.

        Input dataframe df_snpwin consists of 11 columns:
        columns 1, 4, 8 are names of the reference chloroplast genome;
        column 2 and 3 are start and end SNP coordinate;
        columns 5 and 6 are start and end gene coordinate;
        column 7 and 11 are the same as the annotation column (9) from the reference annotation file KC912691.gff3;
        columns 9 and 10 are start and end coordinate of the 1000 bp window in which the respective SNP is located.

        Input dataframe df_nbwin consists of two columns:
        column 1 is the annotation column (9) from the reference annotation file KC912691.gff3 for each gene;
        column 2 represents the number of windows per respective gene from the column 1.
```

```
importing input dataframes
df_snpwin = pd.read_csv(path + 'snps_per_gene_window.bed', sep="\t", header=None)
df_nbwin = pd.read_csv(path + 'number_of_windows_per_gene.csv', sep="\t", header=None)

removing unnecessary columns from df_snpwin
df_snpwin.drop(df_snpwin.columns[[3, 4, 5, 7]], axis = 1, inplace=True)
adding column names to df_snpwin
df_snpwin.columns = ['ref','snp_start','snp_end','attribute1','gene_start','gene_end','attribute2']

removing duplicated attribute columns
df_snpwin_rmdup = df_snpwin.query('attribute1 == attribute2')
df_snpwin_rmdup.drop(['attribute2'], axis=1, inplace=True)

adding column names to df_nbwin
df_nbwin.columns = ['attribute1','nb_windows']
merging df_snpwin and df_nbwin based on their values in attribute1 column
df_norm = pd.merge(df_snpwin_rmdup, df_nbwin, on=['attribute1'], how="outer", indicator=True)

extracting only rows with same values in the "attributte1" column in both dataframes
df_genes_withsnp = df_norm.loc[df_norm['_merge']=='both']
removing the indicator column
df_genes_withsnp.drop(['_merge'], axis=1, inplace=True)

creating a dataframe that contains information about 1000 bp windows without a SNP
df_genes_nosnp = df_norm.loc[df_norm['_merge']=='right_only']
removing the indicator column
df_genes_nosnp.drop(['_merge'], axis=1, inplace=True)

Importing dataframe with coordinates of all genes in the reference chloroplast genome.
```

```
The input dataframe consists of 4 columns: column 1 is the name of the reference chloroplast genome;
columns 2 and 3 are start and end gene coordinate;
column 4 is the annotation column (column 9) from the reference annotation file KC912691.gff3 for each gene.
df_coordinates_all_genes = pd.read_csv(path + 'KC912691_chloroplast.bed', sep="\t", header=None)
adding column names to the input dataframe
df_coordinates_all_genes.columns = ['ref','gene_start','gene_end','attribute1']

merging the dataframe with the information about 1000 bp windows with no SNPs and
dataframe with gene coordinates from all genes from the chloroplast genome
based on their values in the attribute1 column
df_norm_nosnps = pd.merge(df_genes_nosnp, df_coordinates_all_genes, on=['attribute1'],
                                                                      how="outer", indicator=True)
extracting only lines the same value in the attribute1 column in both dataframes
df_norm_nosnps = df_norm_nosnps.loc[df_norm_nosnps['_merge'] == "both"]
removing the unnecessary columns
df_norm_nosnps.drop(['ref_x', 'gene_start_x','gene_end_x','_merge'], axis=1, inplace=True)
ordering the columns of a merged dataframe
df_norm_nosnps.columns = ['snp_start','snp_end','attribute1', 'nb_windows', 'ref', 'gene_start', 'gene_end']
filling all missing values with 0 in the merged dataframe
df_norm_nosnps.fillna(0)

creating dataframe for normalization by merging dataframes with information about 1000 bp windows with and without SNPs
df_norm_final = pd.merge(df_genes_withsnp, df_norm_nosnps, on=['snp_start','snp_end','attribute1','nb_windows','ref','gene_start','gene_end'], how="outer", indicator=True)
splitting the attribute column to get the gene functional group abbreviation
df_norm_final[['first_attribute', 'last_attribute']] = df_norm_final.attribute1.str.split("gene=", expand = True)
df_norm_final[['last_attribute1', 'last_attribute2']] = df_norm_final.last_attribute.str.split(";gene_", expand = True)
removing newly created columns with unnecessary information
df_norm_final.drop(['first_attribute','last_attribute','last_attribute2','_merge'], axis=1, inplace=True)
renaming columns
df_norm_final.rename(columns = {'last_attribute1':'gene'}, inplace = True)
df_norm_final.rename(columns = {'attribute1':'attribute'}, inplace = True)
trimming the values from the "gene" column to get the abbreviation of the gene functional group
df_norm_final['gene_function'] = df_norm_final['gene'].str[:3]
adding gene function for the trnk-UUU gene since it belongs to mat gene family
```

```
df_norm_final.loc[df_norm_final['gene'] == 'trnK-UUU', 'gene_function'] = 'mat'
adding gene function for the photosystem I (PSI) genes
df_norm_final.loc[(df_norm_final['gene'] == 'psaA') |
                  (df_norm_final['gene'] == 'psaB') |
                  (df_norm_final['gene'] == 'psaC') |
                  (df_norm_final['gene'] == 'psaJ') |
                  (df_norm_final['gene'] == 'psaL') |
                  (df_norm_final['gene'] == 'psaF') |
                  (df_norm_final['gene'] == 'psaE') , 'gene_function'] = 'psaI'


df_norm_final = pd.merge(df_norm_final,df1, on=['attribute','gene'], how="outer",
    indicator=True)
df_norm_final = df_norm_final.loc[df_norm_final['_merge']=="both"]
removing gene duplicates
df_norm_final = df_norm_final.drop_duplicates(subset=['gene'], keep= 'first')
normalization (dividing number of SNPs per gene with the total number of windows)
df_norm_final['nb_aSNP_norm'] = df_norm_final['nb_aSNP'] / df_norm_final['nb_windows']
removing unnecessary columns
df_norm_final.drop(['_merge'], axis=1,inplace=True)

adding annotation column for each gene functional group (family)
df_norm_final['families'] = ''
df_norm_final.loc[df_norm_final['gene_function'] == 'atp', 'families'] = 'ATP synthase'
df_norm_final.loc[df_norm_final['gene_function'] == 'ycf', 'families'] = 'hypothetical
    chloroplast ORF'
df_norm_final.loc[df_norm_final['gene_function'] == 'ndh', 'families'] = 'NADH dehydrogenase'
df_norm_final.loc[df_norm_final['gene_function'] == 'pet', 'families'] = 'cytochrome b/f
    complex'
df_norm_final.loc[df_norm_final['gene_function'] == 'psb', 'families'] = 'photosystem II'
df_norm_final.loc[df_norm_final['gene_function'] == 'rpl', 'families'] = 'ribosomal proteins
    (LSU)'
df_norm_final.loc[df_norm_final['gene_function'] == 'rps', 'families'] = 'ribosomal proteins
    (SSU)'
df_norm_final.loc[df_norm_final['gene_function'] == 'rpo', 'families'] = 'RNA polymerase'
df_norm_final.loc[df_norm_final['gene_function'] == 'trn', 'families'] = 'tRNAs'
df_norm_final.loc[df_norm_final['gene_function'] == 'rbc', 'families'] = 'RubisCO large
    subunit'
df_norm_final.loc[df_norm_final['gene_function'] == 'psaI', 'families'] = 'photosystem I'
df_norm_final.loc[df_norm_final['gene_function'] == 'psa', 'families'] = 'photosystem II'
df_norm_final.loc[(df_norm_final['gene_function'] == 'ccs') |
                                  (df_norm_final['gene_function'] == 'cem') |
                                  (df_norm_final['gene_function'] == 'inf'), 'families'] =
                                      'other genes'
df_norm_final.loc[(df_norm_final['gene_function'] == 'mat') |
                  (df_norm_final['gene_function'] == 'clp'), 'families'] = 'clpP, matK'
```

```
adding color column that contains hexadecimal color code for each gene functional group
df_norm_final['color'] = ''
df_norm_final.loc[df_norm_final['families'] == 'ATP synthase', 'color'] = '#CBFFA9'
df_norm_final.loc[df_norm_final['families'] == 'hypothetical chloroplast ORF', 'color'] = '#F1F6F9'
df_norm_final.loc[df_norm_final['families'] == 'NADH dehydrogenase', 'color'] = '#FFE79B'
df_norm_final.loc[df_norm_final['families'] == 'cytochrome b/f complex', 'color'] = '#C8E4B2'
df_norm_final.loc[df_norm_final['families'] == 'photosystem I', 'color'] = 'green'
df_norm_final.loc[df_norm_final['families'] == 'photosystem II', 'color'] = '#8EAC50'
df_norm_final.loc[df_norm_final['families'] == 'ribosomal proteins (LSU)', 'color'] = '#9E6F21'
df_norm_final.loc[df_norm_final['families'] == 'ribosomal proteins (SSU)', 'color'] = '#EEE3CB'
df_norm_final.loc[df_norm_final['families'] == 'RNA polymerase', 'color'] = '#B04759'
df_norm_final.loc[df_norm_final['families'] == 'tRNAs', 'color'] = '#537188'
df_norm_final.loc[df_norm_final['families'] == 'other genes', 'color'] = '#9336B4'
df_norm_final.loc[df_norm_final['families'] == 'clpP, matK', 'color'] = 'orange'
df_norm_final.loc[df_norm_final['families'] == 'RubisCO large subunit', 'color'] = '#B31312'

calculating normalized number of SNPs per each gene functional group
grouped_functions_norm = df_norm_final.groupby(by=['gene_function'])['nb_aSNP'].sum()
grouped_functions_norm_df = grouped_functions_norm.reset_index()
preparing dataframe for plotting for the Fig. 3B
grouped_functions_for_plot = pd.merge(grouped_functions_norm_df, df_norm_final, on=['gene_function'], how="outer", indicator=True)
grouped_functions_for_plot = grouped_functions_for_plot.loc[grouped_functions_for_plot["_merge"]=="both"]
grouped_functions_for_plot = grouped_functions_for_plot.drop_duplicates(subset=["nb_aSNP_x", 'families'], keep="first")

grouped_functions_for_plotf = grouped_functions_for_plot.groupby(by=['families','color'])['nb_aSNP_x'].sum()
grouped_functions_for_plot_df = grouped_functions_for_plotf.reset_index()
------------------------------------------------------------------------------------------------
```

```
-----------------------SNP number distribution per gene function (Fig. 3B)--------------------

mpl.rcParams['font.family'] = 'Arial'

sns.set_theme(style="whitegrid", context="talk")
sns.set_style("whitegrid", {'grid.color': 'lightgray', 'grid.linestyle': '--', 'axes.edgecolor': 'lightgray'})
rcParams['figure.figsize'] = 11.7,8.27
```

```
p = sns.barplot(y="nb_aSNP_x", x="families", data=grouped_functions_for_plot_df,
palette = grouped_functions_for_plot_df['color'],
linewidth = 0.5, edgecolor = "black", ci=None, estimator=sum)

plt.title('SNP distribution per gene function', fontsize=18, font = 'Arial')
p.set_xlabel("Function of genes on chloroplast DNA",fontsize=18, font = 'Arial')
p.set_ylabel("Number of SNPs",fontsize=18, font = 'Arial')
p.set_xticklabels(p.get_xticklabels(),fontsize=16, rotation=90, font = 'Arial')
p.set_yticklabels(p.get_yticks(), size = 16, font = 'Arial')
p.tick_params(axis="y", pad=-5)
p.tick_params(axis="x", pad=-5)
p.bar_label(p.containers[0], fmt= '%0.0f', label_type='center', fontsize = 16, font = 'Arial')

plt.gca().yaxis.set_major_formatter(FuncFormatter(lambda y, _: int(y)))
plt.show()
----------------------------------------------------------------------------------------------
```

```

--------------SNP distribution per 1000 bp window in genes with SNP (Fig. 4A)-----------------

mpl.rcParams['font.family'] = 'Arial'

sns.set_theme(style="whitegrid", context="talk")
sns.set_style("whitegrid", {'grid.color': 'lightgray', 'grid.linestyle': '--',
    'axes.edgecolor': 'lightgray'})
rcParams['figure.figsize'] = 13.7,10.27

p = sns.barplot(y="nb_aSNP_norm", x="gene", data=df_norm_final,
palette = df_norm_final['color'], dodge=False,
linewidth = 0.5, edgecolor = "black", ci=None)

plt.title('SNP distribution per gene', fontsize=18, font = 'Arial')
p.set_xlabel("Chloroplast genes", fontsize=18, font = 'Arial')
p.set_ylabel("Number of SNPs / 1000 bp gene length", fontsize=18, font = 'Arial')
p.set_xticklabels(p.get_xticklabels(), fontsize=16, rotation=90, font = 'Arial')
p.set_yticklabels(p.get_yticks(), size = 16, font = 'Arial')
p.tick_params(axis="y", pad=-5)
p.tick_params(axis="x", pad=-5)
p.bar_label(p.containers[0], fmt= '%0.1f', label_type='edge', fontsize = 7, font = 'Arial')
p.relim()
p.autoscale_view()
p.margins(x=0.002)


plt.gca().yaxis.set_major_formatter(FuncFormatter(lambda y, _: int(y)))
plt.show()
```

```
----------------------------------------------------------------------------------------------
```

```

------SNP distribution per 1000 bp window 10 genes with the highest SNP number (Fig. 4B)------

mpl.rcParams['font.family'] = 'Arial'

sns.set_theme(style="whitegrid", context="talk")
sns.set_style("whitegrid", {'grid.color': 'lightgray', 'grid.linestyle': '--',
    'axes.edgecolor': 'lightgray'})
rcParams['figure.figsize'] = 13.7,10.27

df_norm_final_max = df_norm_final.sort_values(by=['nb_aSNP_norm'], ascending=False).head(10)

p = sns.barplot(y="nb_aSNP_norm", x="gene", data=df_norm_final_max,
palette = df_norm_final_max['color'], dodge=False,
linewidth = 0.5, edgecolor = "black", ci=None)

plt.title('SNP distribution per gene', fontsize=18, font = 'Arial')
p.set_xlabel("Chloroplast genes", fontsize=18, font = 'Arial')
p.set_ylabel("Number of SNPs / 1000 bp gene length", fontsize=18, font = 'Arial')
p.set_xticklabels(p.get_xticklabels(), fontsize=16, rotation=90, font = 'Arial')
p.set_yticklabels(p.get_yticks(), size = 16, font = 'Arial')
p.tick_params(axis="y", pad=-5)
p.tick_params(axis="x", pad=-5)
p.bar_label(p.containers[0], fmt= '%0.1f', label_type='center', fontsize = 16, font = 'Arial')
p.relim()
p.autoscale_view()
p.margins(x=0.002)


plt.gca().yaxis.set_major_formatter(FuncFormatter(lambda y, _: int(y)))
plt.show()
----------------------------------------------------------------------------------------------
```

```
---------------------05_generating_upset_and_circos_plotting_dataframes.py--------------------

Purpose: Generating a dataframe for the upset plot (Fig. S1; Fig. 2A);
         Generating a dataframe with chloroplast gene coordinates
         for the part of the circos plot (Fig. 3A)
```

```
--------------------------creating plotting dataframe for Fig. S1-----------------------------

importing necessary modules
import pandas as pd
import numpy as np

```

```
setting input file directory path
vcf_path = '/path/to/merged/vcf/file/with/SNPs/from/all/isolates/'

removing all of the lines before the line of a vcf file that starts with #CHROM
with open(vcf_path + 'merged_vcf_file.vcf') as infile:
    for n,row in enumerate(infile):
        if row.startswith('#CHROM'):
            break

importing the dataframe (merged vcf file) starting from the line that has '#CHROM' at the beginning
df = pd.read_csv(vcf_path + 'merged_vcf_file.vcf', sep="\t", skiprows=n)
removing unnecessary columns from the imported vcf file
df.drop(['#CHROM', 'ID', 'REF', 'ALT', 'QUAL', 'FILTER', 'INFO', 'FORMAT'], axis=1, inplace=True)
making a list of column names
list_of_column_names = df.columns.to_list()

full_column_names = []

extracting the accession name of each column name that has more than 8 characters

def extracting_the_accession_name():
    for name in list_of_column_names:
        print(name)
        if len(name) > 8:
            full_column_names.append(name[:8])
        else:
            pass

inserting 'SNP' on the first position of the list of column names
full_column_names.insert(0, 'SNP')

df.columns = full_column_names

replacing all dots with nan values and storing it into new df1 dataframe
df1 = df.replace('.', np.nan)

replacing all not nan values from the imported dataframe (df1) with 1 and storing it into new dataframe
df2 = df1.notnull().astype('int')

removing the first column of the df2 since SNP positions were replaced with 1 with the command above
df2.drop('SNP', axis=1, inplace=True)

```

```
concatenating the first column of df1 to the columns of df2 and storing it to a new dataframe (df3)
df3 = pd.concat([df1['SNP'], df2], axis=1)

exporting the dataframe for the upset plot (Fig. S1)
df3.to_csv(vcf_path + 'upset_df.csv', sep='\t', index=False)
```

```

---------------------------creating plotting dataframe for Fig. 2A----------------------------

The final dataframe for each location will have two columns:
col1---SNP positions;
col2---0/1 values for each position for all isolates from the corresponding location
depending on the SNP presence/absence on specific location


make a list with isolate IDs from the corresponding location for each location
Gottingen = ['KP210080']
Zaisenhausen_Wackershofen = ['KP210099']
Weipertshofen = ['KP210124']
Schwabisch_Hall = ['KP210160','KP210161','KP210169','KP210170','KP210171']
Reutlingen = ['KP210195','KP210196','KP210197']

df_Gottingen = df3[df3.columns.intersection(Gottingen)]
df_Zaisenhausen_Wackershofen = df3[df3.columns.intersection(Zaisenhausen_Wackershofen)]
df_Weipertshofen = df3[df3.columns.intersection(Weipertshofen)]
df_Schwabisch_Hall = df3[df3.columns.intersection(Schwabisch_Hall)]
df_Reutlingen = df3[df3.columns.intersection(Reutlingen)]
```

```
merging of values from all columns of dataframes for every sampling site into a single column
df_Gottingen['Gottingen'] = df_Gottingen.sum(axis=1, numeric_only=True)
df_Zaisenhausen_Wackershofen['Zaisenhausen/Wackershofen'] =
    df_Zaisenhausen_Wackershofen.sum(axis=1, numeric_only=True)
df_Weipertshofen['Weipertshofen'] = df_Weipertshofen.sum(axis=1, numeric_only=True)
df_Schwabisch_Hall['Schwabisch Hall'] = df_Schwabisch_Hall.sum(axis=1, numeric_only=True)
df_Reutlingen['Reutlingen'] = df_Reutlingen.sum(axis=1, numeric_only=True)

df_Gottingen.drop(df_Gottingen.columns.difference(['Gottingen']), 1, inplace=True)
df_Zaisenhausen_Wackershofen.drop(
    df_Zaisenhausen_Wackershofen.columns.difference(['Zaisenhausen/Wackershofen']), 1, inplace=True)
df_Weipertshofen.drop(df_Weipertshofen.columns.difference(['Weipertshofen']), 1, inplace=True)
df_Schwabisch_Hall.drop(df_Schwabisch_Hall.columns.difference(['Schwabisch Hall']), 1, inplace=True)
df_Reutlingen.drop(df_Reutlingen.columns.difference(['Reutlingen']), 1, inplace=True)

df_Gottingen[df_Gottingen > 1] = 1
```

```
df_Zaisenhausen_Wackershofen[df_Zaisenhausen_Wackershofen > 1] = 1
df_Weipertshofen[df_Weipertshofen > 1] = 1
df_Schwabisch_Hall[df_Schwabisch_Hall > 1] = 1
df_Reutlingen[df_Reutlingen > 1] = 1
```

```
merge final dataframes of every sampling site into a single dataframe
df4 = pd.concat([df3['SNP'],
                 df_Gottingen,
                 df_Zaisenhausen_Wackershofen,
                 df_Weipertshofen,
                 df_Schwabisch_Hall,
                 df_Reutlingen], axis=1)

exporting the dataframe for upset plot
df4.to_csv(vcf_path + 'upset_df_locations.csv', sep='\t', index=False)
---------------------------------------------------------------------------------------------
```

```
----------------------------creating plotting dataframe for Fig. 3A---------------------------

importing the bed file that is output of the 03_SNP_per_gene.sh script
df_circos_1 = pd.read_csv(vcf_path + 'KC912691_genes.bed', sep="\t")

adding the gene function column to df_circos_1
df_circos_1['gene_function'] = df_circos_1['gene'].str[:3]
adding gene function for the trnk-UUU gene since it belongs to mat gene family
df_circos_1.loc[df_circos_1['gene'] == 'trnK-UUU', 'gene_function'] = 'mat'
adding gene function for the photosystem I (PSI) genes
df_circos_1.loc[(df_circos_1['gene'] == 'psaA') |
                (df_circos_1['gene'] == 'psaB') |
                (df_circos_1['gene'] == 'psaC') |
                (df_circos_1['gene'] == 'psaJ') |
                (df_circos_1['gene'] == 'psaL') |
                (df_circos_1['gene'] == 'psaF') |
                (df_circos_1['gene'] == 'psaE') , 'gene_function'] = 'psaI'

adding annotation column for each gene functional group (family)
df_circos_1['families'] = ''
df_circos_1.loc[df_circos_1['gene_function'] == 'atp', 'families'] = 'ATP synthase'
df_circos_1.loc[df_circos_1['gene_function'] == 'ycf', 'families'] = 'hypothetical chloroplast ORF'
df_circos_1.loc[df_circos_1['gene_function'] == 'ndh', 'families'] = 'NADH dehydrogenase'
df_circos_1.loc[df_circos_1['gene_function'] == 'pet', 'families'] = 'cytochrome b/f complex'
df_circos_1.loc[df_circos_1['gene_function'] == 'psb', 'families'] = 'photosystem II'
df_circos_1.loc[df_circos_1['gene_function'] == 'rpl', 'families'] = 'ribosomal proteins (LSU)'
df_circos_1.loc[df_circos_1['gene_function'] == 'rps', 'families'] = 'ribosomal proteins (SSU)'
```

```
df_circos_1.loc[df_circos_1['gene_function'] == 'rpo', 'families'] = 'RNA polymerase'
df_circos_1.loc[df_circos_1['gene_function'] == 'trn', 'families'] = 'tRNAs'
df_circos_1.loc[df_circos_1['gene_function'] == 'psaI', 'families'] = 'photosystem I'
df_circos_1.loc[df_circos_1['gene_function'] == 'psa', 'families'] = 'photosystem II'
df_circos_1.loc[df_circos_1['gene_function'] == 'rbc', 'families'] = 'RubisCO large subunit'
df_circos_1.loc[(df_circos_1['gene_function'] == 'ccs') |
                (df_circos_1['gene_function'] == 'cem') |
                (df_circos_1['gene_function'] == 'inf'), 'families'] = 'other genes'
df_circos_1.loc[(df_circos_1['gene_function'] == 'mat') |
                (df_circos_1['gene_function'] == 'clp'), 'families'] = 'clpP, matK'

adding color column that contains hexadecimal color code for each gene functional group
df_circos_1['color'] = ''
df_circos_1.loc[df_circos_1['families'] == 'ATP synthase', 'color'] = '#CBFFA9'
df_circos_1.loc[df_circos_1['families'] == 'hypothetical chloroplast ORF', 'color'] =
    '#F1F6F9'
df_circos_1.loc[df_circos_1['families'] == 'NADH dehydrogenase', 'color'] = '#FFE79B'
df_circos_1.loc[df_circos_1['families'] == 'cytochrome b/f complex', 'color'] = '#C8E4B2'
df_circos_1.loc[df_circos_1['families'] == 'photosystem I', 'color'] = 'green'
df_circos_1.loc[df_circos_1['families'] == 'photosystem II', 'color'] = '#8EAC50'
df_circos_1.loc[df_circos_1['families'] == 'ribosomal proteins (LSU)', 'color'] = '#9E6F21'
df_circos_1.loc[df_circos_1['families'] == 'ribosomal proteins (SSU)', 'color'] = '#EEE3CB'
df_circos_1.loc[df_circos_1['families'] == 'RNA polymerase', 'color'] = '#B04759'
df_circos_1.loc[df_circos_1['families'] == 'tRNAs', 'color'] = '#537188'
df_circos_1.loc[df_circos_1['families'] == 'other genes', 'color'] = '#9336B4'
df_circos_1.loc[df_circos_1['families'] == 'clpP, matK', 'color'] = 'orange'
df_circos_1.loc[df_circos_1['families'] == 'RubisCO large subunit', 'color'] = '#B31312'

removing unnecessary columns
df_circos_1.drop(['gene_function'], axis=1, inplace=True)
renaming columns
df_circos_1.rename(columns={'families': 'gene_function'}, inplace=True)
exporting dataframe
df_circos_1.to_csv(vcf_path + 'circos_genes.bed', sep="\t", index=False)
--------------------------------------------------------------------------------------------
```

```
--------------------------------06_upset_and_circos_plots.R------------------------------------

Purpose: R script that takes as input dataframes generated in scripts:
         02_variant_annotation.py
         05_generating_upset_and_circos_plotting_dataframes.py
         and creates plots for Fig. 2A, Fig. S1 and Fig. S3A
```

```
importing necessary libraries
library(UpSetR)
library(ggplot2)

```

```
importing input dataframes
variants_per_isolate <- read.csv('upset_df.csv', sep = '\t')
variants_per_sampling_site <- read.csv('upset_df_locations.csv', sep = '\t')


----------------------------------------plotting Fig. S1--------------------------------------

upset(variants_per_isolate, nsets = 11, point.size = 4, line.size = 1.5,
      mainbar.y.label = "Number of shared SNPs between isolates", sets.x.label = "SNP number per isolate",
      sets = c('KP210080', 'KP210099', 'KP210124',
               'KP210160', 'KP210161', 'KP210169',
               'KP210170', 'KP210171', 'KP210195', 'KP210196', 'KP210197'),
      nintersects = NA,
      keep.order = T,
      main.bar.color = '#aad400ff',
      text.scale = c(1.5, 1.6, 1.3, 1, 2, 1.5),
      sets.bar.color=c("#008000ff",
                       "#37c837ff",
                       "#2aff80ff",
                       "#00bd7dff","#00bd7dff","#00bd7dff","#00bd7dff","#00bd7dff",
                       "#7fff2aff","#7fff2aff","#7fff2aff"))
----------------------------------------------------------------------------------------------

----------------------------------------plotting Fig. 2A--------------------------------------

upset(variants_per_sampling_site, nsets = 5, point.size = 4, line.size = 1.5,
      mainbar.y.label = "Number of SNP intersections between locations",
      sets.x.label = "SNP number per location",
      sets = c('Gottingen', 'Zaisenhausen.Wackershofen',
               'Weipertshofen','Schwabisch.Hall', 'Reutlingen'),
      nintersects = NA,
      keep.order = T,
      main.bar.color = '#aad400ff',
      text.scale = c(1.5, 1.6, 1.3, 1, 2, 1.5),
      sets.bar.color=c("#008000ff",
                       "#37c837ff",
                       "#2aff80ff",
                       "#00bd7dff",
                       "#7fff2aff"))
----------------------------------------------------------------------------------------------
```

```

----------------------------------------plotting Fig. 3A--------------------------------------

installing packages and libraries
```

```
install.packages("circlize")
devtools::install_github("jokergoo/circlize")

if (!require("BiocManager", quietly = TRUE))
  install.packages("BiocManager")

BiocManager::install("ComplexHeatmap")

library(circlize)
library(ggsignif)
library(ComplexHeatmap)


--------------------------The structure of the input dataframes:------------------------------
df dataframe contains start and end coordinate of the chloroplast genome:
chr start   end
KC912691.1  0   114843

variants dataframe is generated by 02_variant_annotation.py script:
chr position    start   end type    color
KC912691.1  4548    4548    4549    synonymous_variant  #92C5DE
KC912691.1  9456    9456    9457    synonymous_variant  #92C5DE

genes dataframe is generated by 05_generating_upset_and_circos_plotting_dataframes.py script:
chr start   end gene    gene_function   color
KC912691.1  75  1136    psbA    photosystem II  #8EAC50
KC912691.1  1358    3931    trnK-UUU    clpP, matK  orange
----------------------------------------------------------------------------------------------
```

```
importing dataframes
df <- read.csv("circos_genome.bed", sep="\t")
variants <- read.csv("circos_snps.bed", sep="\t")
genes <- read.csv("circos_genes.bed", sep="\t")

removing underscore from the column type of the variants dataframe
variants$type<-gsub("_"," ",as.character(variants$type))

generating circos plot
circos.clear()

circos.par("track.height"=0.2, gap.degree=0, start.degree = 190, cell.padding=c(0, 0, 0, 0))




p <- circos.initialize(factors = c("KC912691.1"),
```

```
                       xlim = matrix(c(rep(0, 1), df$end), ncol=2),
                       circos.track(ylim = c(0, 1)),
                       set_track_gap(mm_h(0.0)))

circos.track(ylim=c(0, 1), panel.fun=function(x, y) {
  chr=CELL_META$sector.index
  xlim=CELL_META$xlim
  ylim=CELL_META$ylim
  circos.text(mean(xlim), mean(ylim), chr, cex=0.5, col="white",
              facing="outside", niceFacing=TRUE)
},bg.col= c("white"), bg.border=F, track.height=0.03)

brk <- c(1.16485)*10^5
circos.track(track.index = get.current.track.index(), panel.fun=function(x, y) {
  circos.axis(h="top", major.at=brk, labels=round(brk/10^5, 1), labels.cex=0.4,
              col="white", labels.col="white", lwd=0.7, labels.facing="clockwise")
}, bg.border=F)

for (j in 1:nrow(variants)){
  circos.lines(x=round(variants$position[j]),
               y=1,
               type="h",
               baseline = "bottom",
               col = "black", border = "black")

  circos.points(x=round(variants$position[j]),
                     y=1,
                     col=variants$color[j],
                     cex = 0.7,
                     pch = 19)}


circos.genomicTrack(genes, ylim = c(0, 1),
                    panel.fun = function(region, value, ...) {
                      circos.genomicRect(region, value, col = genes$color,
                                         border = F)

                      cell.xlim = get.cell.meta.data("cell.xlim")
                    }, track.height=0.04, bg.border=F)

circos.genomicTrack(df, ylim = c(0, 1),
                    panel.fun = function(region, value, ...) {
                      circos.genomicRect(region, value, col = "black",
                                         border = F)

                      cell.xlim = get.cell.meta.data("cell.xlim")
```

```
                    }, track.height=0.005, bg.border=F)

circos.genomicLabels(genes,
                     labels = NULL,
                     labels.column = "gene",
                     facing = "reverse.clockwise",
                     niceFacing = TRUE,
                     col = "black",
                     cex = 0.5,
                     padding = 0.4,
                     line_col = "black",
                     line_lwd = 0.7,
                     line_lty = 1,
                     side = c("inside"))

generating legend for the circos plot
nms_lgd = c("photosystem I","photosystem II","cytochrome b/f complex", "ATP synthase",
            "NADH dehydrogenase","RubisCO large subunit","RNA polymerase",
            "ribosomal proteins (LSU)", "ribosomal proteins (SSU)",
            "tRNAs", "clpP, matK", "other genes", "hypothetical chloroplast ORF")

clr_lgd = c("green", "#8EAC50", "#C8E4B2", "#CBFFA9",
            "#FFE79B","#B31312","#B04759", "#9E6F21", "#EEE3CB",
            "#537188","orange", "#9336B4", "#F1F6F9")

lgd_1 = Legend(title = "Gene function", at = c(nms_lgd), title_position = "topleft",
               legend_gp = gpar(fill = clr_lgd))

lgd_2 = Legend(title = "SNP mutation", at = unique(variants$type), title_position = "topleft",
               legend_gp = gpar(fill = unique(variants$color)),  type = "p", pch = 21)


lgd_list_vertical = packLegend(lgd_1, lgd_2)
draw(lgd_list_vertical)
---------------------------------------------------------------------------------------------
```
