## Supplementary figures and images for "Sequencing of historical plastid genomes reveal exceptional genetic diversity in early domesticated rye plants"

### FigS1.pdf

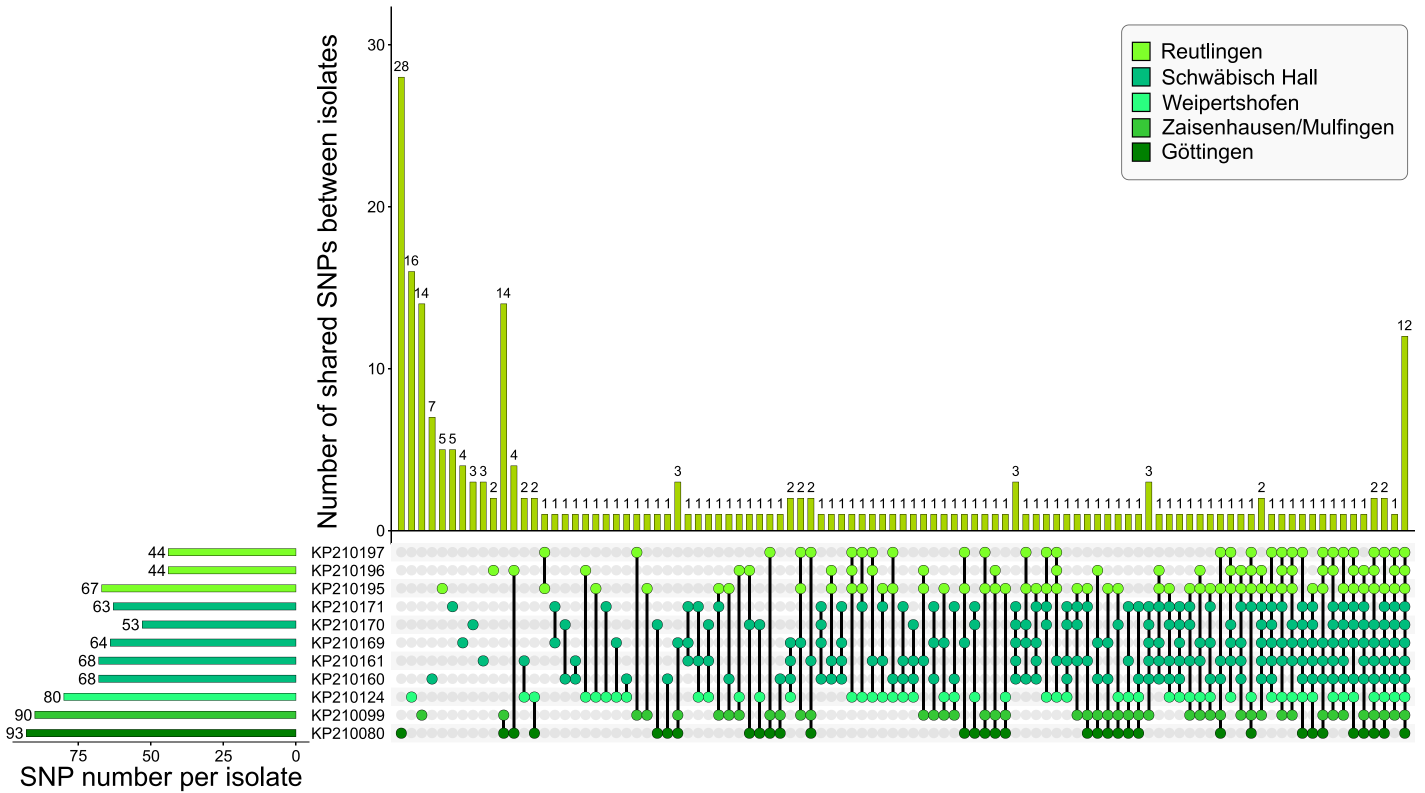
